# Decoding functional cell–cell communication events by multi-view graph learning on spatial transcriptomics

**DOI:** 10.1101/2022.06.22.496105

**Authors:** Haochen Li, Tianxing Ma, Minsheng Hao, Wenbo Guo, Jin Gu, Lei Wei, Xuegong Zhang

## Abstract

Cell–cell communication events (CEs) are mediated by multiple ligand–receptor pairs. Usually only a particular subset of CEs directly works for a specific downstream response in a particular microenvironment. We name them as functional communication events (FCEs) of the target responses. Decoding the FCE-target gene relations is important for understanding the machanisms of many biological processes, but has been intractable due to the mixing of multiple factors and the lack of direct observations. We developed a method HoloNet for decoding FCEs using spatial transcriptomic data by integrating ligand–receptor pairs, cell-type spatial distribution and downstream gene expression into a deep learning model. We modeled CEs as a multiview network, developed an attention-based graph learning method to train the model for generating target gene expression with the CE networks, and decoded the FCEs for specific downstream genes by interpreting the trained model. We applied HoloNet on three Visium datasets of breast cancer or liver cancer. It revealed the communication landscapes in tumor microenvironments, and uncovered how various ligand–receptor signals and cell types affect specific biological processes. We also validated the stability of HoloNet in a Slideseq-v2 dataset. The experiments showed that HoloNet is a powerful tool on spatial transcriptomic data to help revealing specific cell–cell communications in a microenvironment that shape cellular phenotypes.

## Introduction

Cell–cell communication is of vital importance to the organization and maintenance of multicellular organisms as well as the occurrence of diseases (Armingol et al., 2020; Bich et al., 2019; Bonnans et al., 2014). A cell– cell communication event (CE) occurs when sender cells release ligand molecules to the microenvironment and receiver cells sense the ligands by receptors. CEs form a complex intercellular signaling network that participates in the regulation of various genes. Abnormal CEs can trigger abnormal cellular phenotypes and may lead to the occurrence of many diseases (Niethamer et al., 2020; Zepp et al., 2017).

A comprehensive description of a CE should include the sender cell, the receiver cell, the ligand–receptor pair, and its downstream responses (**Figure 1A**). It is worth noting that downstream responses are usually induced by only a particular subset of all possible CEs. We call such CEs in the subset as functional communication events (FCEs) for the downstream responses. FCEs refers specifically to CEs that involve specific biological processes and eliminate accidental or irrelevant CEs, and thus can help better understand the role of cell–cell communication in shaping cellular phenotypes (Harney et al., 2015; Selvey et al., 2004) as well as develop possible disease interventions (Liu et al., 2020; Roy et al., 2018). Decoding FCEs should answer the following questions: which cell types act as senders and receivers, where they are, and which ligand–receptor pairs they employ to induce what kind of downstream responses. The answers to all these questions will compose the “holographic” view of the networks of FCEs.

**Figure 1.**
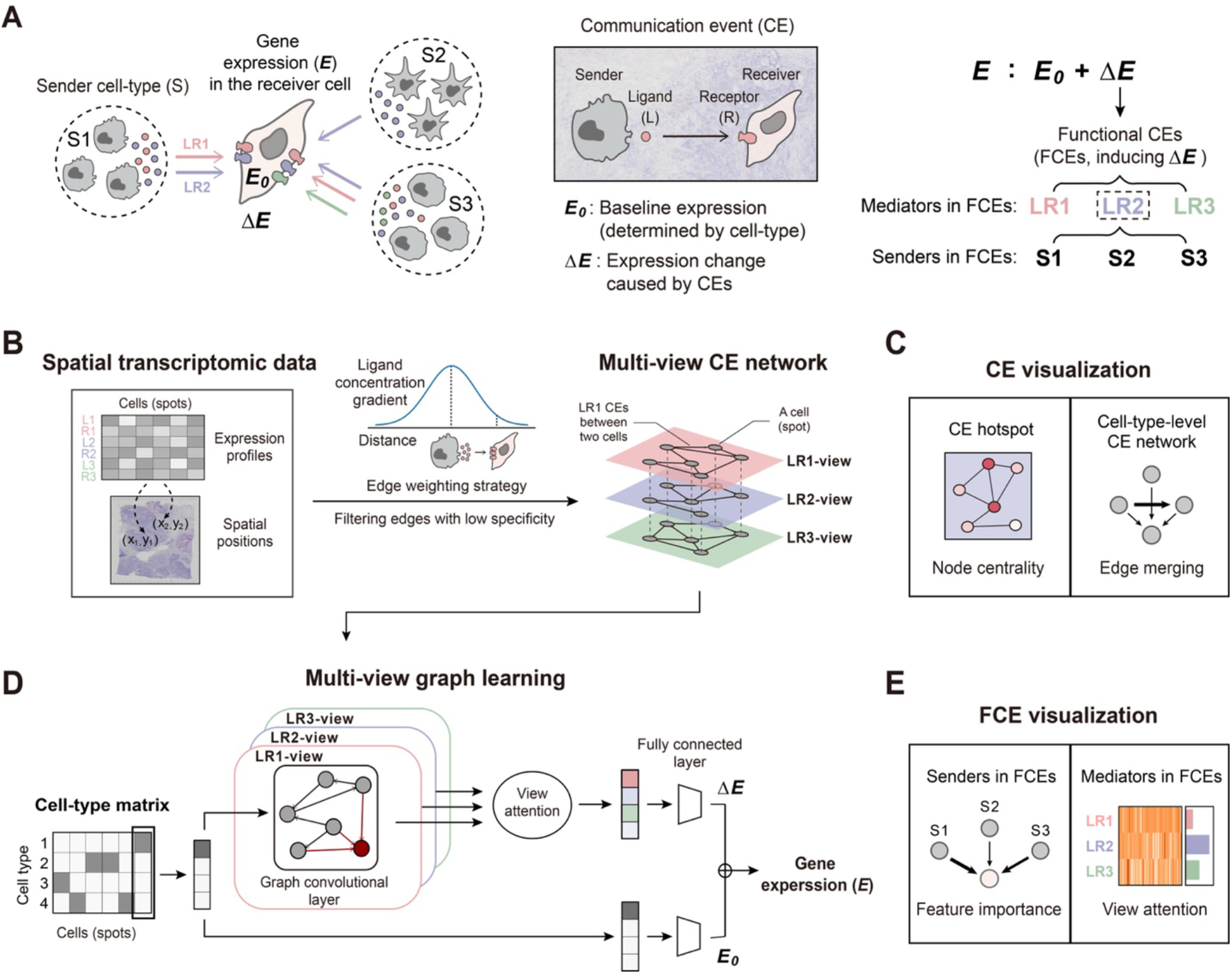
The overall workflow of HoloNet. **(A)** Definition of CEs and FCEs. **(B)** Constructing multi-view CE networks using spatial transcriptomic data. **(C)** HoloNet outputs the visualization results of CEs based on the multi-view CE network. HoloNet provides CE hotspots according to the node centralities in the CE network, and provides cell-type-level CE networks by merging edges from one cell type to another cell type. **(D)** Generating specific gene expressions via multi-view graph learning on the constructed multi-view CE network. First, HoloNet propagates the cell-type information over the multi-view CE network via the graph convolutional layer in each view. Then, HoloNet integrates the embeddings of nodes from each view with attention mechanisms. Finally, HoloNet estimates **Δ*E*** and ***E_0_*** respectively from integrated embeddings and cell-type matrix through a fully connected layer. The generated expressions are the sum of ***E_0_*** and ***ΔE***. **(E)** HoloNet outputs visualization results of FCEs by interpreting the multi-view graph learning model. HoloNet identifies ligand–receptor pairs as the core mediators, and cell types as the core senders of FCEs for specific genes.

Current molecular profiling technologies provide bulk or single-cell gene expression profiles which include major ligand and receptor genes, cell-type-specific marker genes, and possible downstream response genes. Recent spatial molecular profiling technologies such as spatial transcriptomic sequencing (Salmén et al., 2018; Ståhl et al., 2016) and SeqFISH (Eng et al., 2019) can provide additional information about cell positions. Some data-driven methods have been developed to study CEs based on these technologies (Browaeys et al., 2020; Cang and Nie, 2020; Fischer et al., 2021; Hou et al., 2020; Hu et al., 2021; Jin et al., 2021; Li et al., 2020; Ramilowski et al., 2015; Ren et al., 2020). For example, NicheNet (Browaeys et al., 2020) and MESSI (Li et al., 2020) were built to characterize the relationship between ligand–receptor pairs and downstream responses, while SVCA (Arnol et al., 2019) and NCEM (Fischer et al., 2021) focused on the sender–receiver cell dependencies as well as the related phenotypes. However, none of the exsiting methods have focused on FCEs: They only revealed parts of the core elements of FCEs. A method for systematically decoding FCEs is still lacking, which limits the exploration on the detailed roles of various cell types in different biological processes and the transmission process of various ligand–receptor signals in tissues.

In this work, we developed a computational method named HoloNet to decode FCEs using spatial transcriptomic data. We modeled CEs in spatial data as a multi-view network using the ligand and receptor expression profiles, developed a graph neural network model to generate the expressions of specific genes, and then interpreted the trained neural network to decode the full picture of FCEs. HoloNet provides a general workflow to characterize the communication landscapes in spatial data based on the multi-view CE network, and can identify cell types serving as major senders and ligand–receptor pairs serving as core mediators in FCEs for specific downstream genes. We applied HoloNet on three datasets obtained with the 10x Visium technology (Wu et al., 2021a, 2021b), and a dataset obtained with Slideseq-v2 (Stickels et al., 2021). The results provided new insights into how the microenvironment shapes cellular phenotypes in the tissues. HoloNet is available as a Python package at https://github.com/lhc17/HoloNet free for academic use.

## Results

### Overview of HoloNet

HoloNet accepts spatial transcriptomic data, cell-type labels and a list of ligand–receptor gene pairs as inputs. The required spatial data should contain high-dimensional gene expression profiles and spatial positions of cells (**Figure 1B**). Users can provide either continuous cell-type percentages derived from deconvolution methods such as Seurat (Stuart et al., 2019) for spot-based datasets or categorical cell-type labels for datasets with single-cell resolution. We used the paired ligand–receptor gene lists from CellChatDB (Jin et al., 2021) as the default setting, but users can also provide gene lists from other databases.

HoloNet constructs a multi-view CE network using the spatial data and paired ligand–receptor gene lists. As shown in **Figure 1A**, in a CE, ligand molecules are released by the sender cell, diffuse over a distance within the tissue, and then bind to receptors expressed by the receiver cell. As cells communicate with others via multiple ligand–receptor pairs, CEs can be naturally modeled as a multi-view network. Each view in this network is a directed graph representing CEs mediated by one ligand–receptor pair with their own weighted edges. Each node represents a single cell or a spot in the measured tissue, and nodes across the multiple views are aligned. A directed edge in any view connects a sender cell to its corresponding receiver cell. An edge weight represents the strength of a CE mediated by the ligand–receptor pair between the sender and receiver cell. For each view, HoloNet calculates the weight of each edge by a function of three factors: the expression of the ligand in the sender cell, the expression of the receptor in the receiver cell, and the physical distance between the sender and receiver cell (**Figure 1B**, **Methods**). Besides, communication mediated by secretory ligands can occur across the whole tissue, while other communication is limited to adjacent cells. HoloNet models these different properties with different parameter settings to illustrate cell–cell communication more accurately (**Figure S2A, Methods**). HoloNet retains edges with high confidence evaluated via a permutation test (**Figure S1A, Methods**).

HoloNet provides multiple approaches to analyze and visualize the multi-view CE network (**Figure 1C**). For any ligand–receptor pair, HoloNet computes centrality measures in the corresponding view to detect cells (or spots) with high communication activation as CE hotspots. HoloNet also constructs a cell-type-level CE network for each view which represents the general CE strengths from one cell type to another cell type. To explore the relationship between different ligand–receptor pairs, HoloNet quantifies similarities between different network views based on their local features and performs the hierarchical clustering method to cluster ligand–receptor pairs (**Methods**).

HoloNet constructs the FCE network based on the constructed multi-view CE network. We regard a CE as an FCE if the CE alters the expression of the gene of interest in the receiver cell. To decode FCEs, we decomposed the expression ***E*** of the target gene in all single cells or spots into its baseline expression ***E_0_*** determined by its cell type and the expression change **Δ*E*** caused by CEs triggered by multiple cell types and mediated by multiple ligand–receptor pairs (**Figure 1A, Methods**). We employ multi-view graph neural networks (GNNs) to generate the expression profile of the target genes, and interpret the trained model to decode FCEs. **Figure 1D** illustrates the procedure. We first obtained a matrix representing the cell type of each cell by one-hot encoding or the cell type percentages in each spot. Then, a GNN is constructed for each view by adopting the CE network of this view as the adjacency matrix and assigning each column of the cell-type matrix to the corresponding node in each graph as features. Embeddings of nodes of each view are integrated inspired by the attention mechanism (Fu et al., 2022; Khan and Blumenstock, 2019). The integrated results are fed into a fully connected layer to estimate **Δ*E***. Another fully connected layer is constructed to estimate ***E*_0_** from the cell-type matrix. The whole model is trained to minimize the difference between ***E*** and the sum of ***E*_0_** and **Δ*E*** (**Figure 1D, Methods**).

After training, we interpret the attention weights of HoloNet to indicate which views in the multi-view network contribute more to the expression of a target gene. The ligand–receptor pairs corresponding to these views are regarded as the core mediators of FCEs regulating the target gene expression. All attention weights as well as their mean values across all trained models are visualized as the FCE mediator plot. We can further identify the major sender and receiver cell types in FCEs based on the graph convolutional layer to construct cell-type-level FCE connectivity networks (**Figures 1E and S1B, Methods**). To ensure the reliability of the findings, we repeated the training procedure for each target gene and integrated all the model-generated gene expressions and model interpretation results.

We applied HoloNet on four datasets, including two human breast cancer Visium spatial transcriptomic datasets, one human liver cancer Visium spatial transcriptomic dataset and one mouse hippocampus Slideseq-v2 dataset. One of the breast cancer dataset is from the 10x Genomics website (**Methods**), and others are from publications (Stickels et al., 2021; Wu et al., 2021a, 2021b).

### Chatacterizing cell–cell communication events in breast cancer

We first applied HoloNet on the breast cancer Visium dataset from the 10x Genomics website. The dataset is derived from an invasive ductal carcinoma breast tissue sample and it profiled the expression of 24,923 genes in 3,798 spots. We calculated the cell-type percentage of each spot using the deconvolution method in Seurat (Stuart et al., 2019) (**Methods**). The cell types include stromal cells, immunocytes, and tumor cells which are further divided into four cell types: basal-like-1 cells, basal-like-2 cells, luminal-AR cells and mesenchymal cells (Karaayvaz et al., 2018; Lehmann et al., 2016) (**Figures 2A and 2B**).

**Figure 2.**
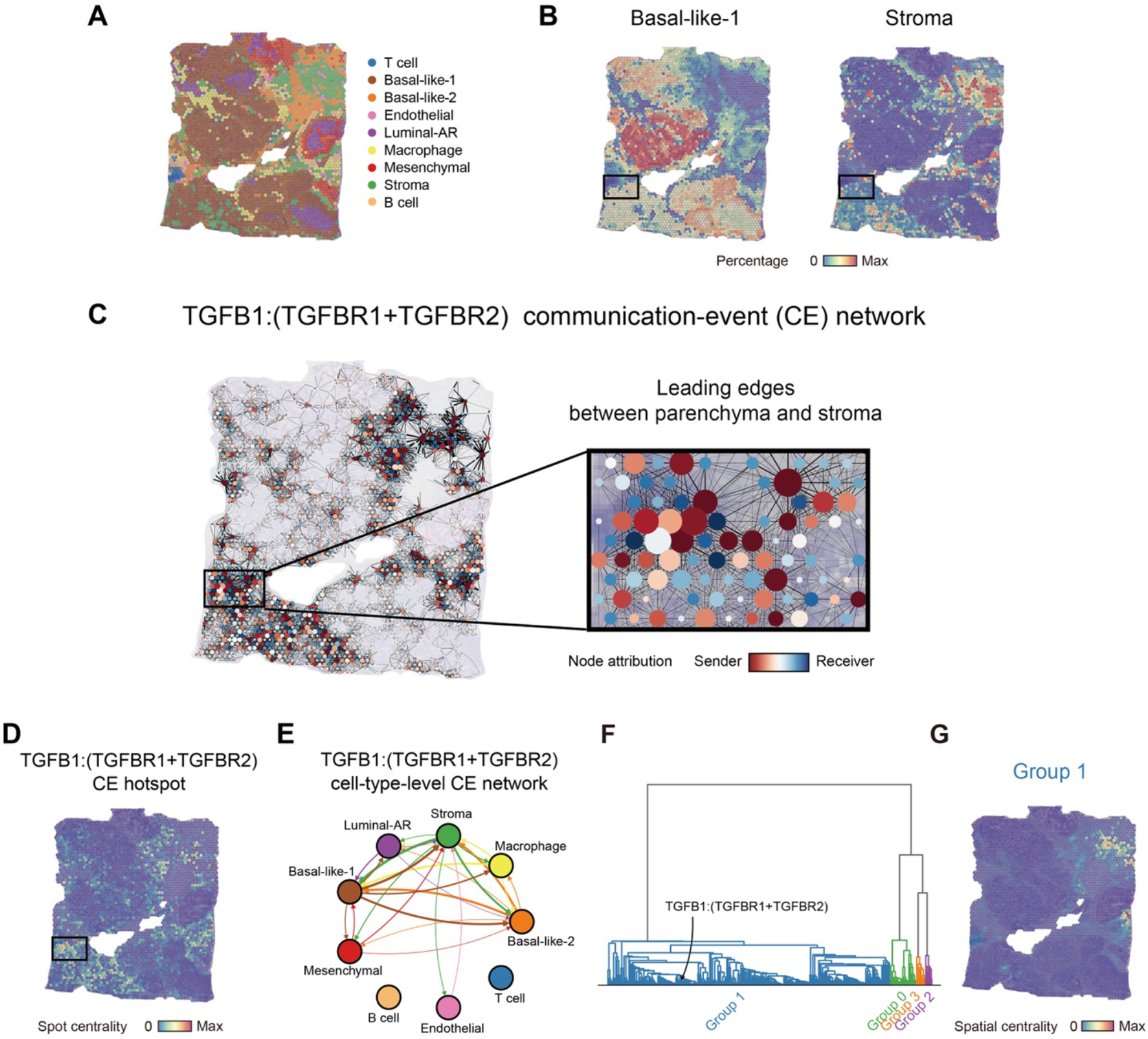
Characterizing cell–cell communication events of a breast cancer Visium dataset by HoloNet. **(A)** The cell type with the highest percentage in each spot. Each dot printed on the slide represents a single invisible capture spot with a diameter of 55 μm. **(B)** The percentages of basal-like-1 and stromal cells in each spot. **(C)** The constructed single-view CE network based on the TGFB1:(TGFBR1+TGFBR2) pair. The size of a node indicates the degree centralities of the node in the network, and the color indicates the sender-receiver attribute of the node. The sender-receiver attribute is the result of subtracting the in-degree of a node from its out-degree. The redder the color of nodes, the more inclined the nodes are to act as a sender, and the bluer on the contrary. The thickness of an edge indicates the strength of CEs between cells linked by it. Nodes with too small size and edges with too low thickness are not displayed. Nodes are mapped to the spatial positions of their corresponding spots in the hematoxylin and eosin (H&E) stained histology image. **(D)** CE hotspot plot describes regions with active TGFB1:(TGFBR1+TGFBR2) signals. For (C), (D) and (E), the leading edge between parenchyma and stroma is highlighted. **(E)** Cell-type-level CE network plot describes the TGFB1:(TGFBR1+TGFBR2) communication strengths between cell types. The thickness of an edge reflects the summed weight of all edges in the CE network between the pair of cell types. **(F)** Dendrogram for hierarchically clustering all ligand–receptor pairs. Ligand–receptor pairs were clustered into four groups based on the eigenvector centrality vectors. **(G)** The general CE hotspot of ligand-receptor Group 1. All hotspots of members in a group are superimposed to generate the general hotspot.

We constructed the multi-view CE network in the dataset by HoloNet. The multi-view CE network connects the 3,798 spots in 325 views, and each view corresponds to a ligand–receptor pair. Here we took the ligand TGF-β1 (TGFB1) and the corresponding receptor complex composed of TGF-β receptor 1 and 2 (TGFBR1 and TGFBR2) as an example (**Figure S2B**). This ligand–receptor pair participates in many critical biological processes in many cell types, such as inducing epithelial-mesenchymal transition in tumor cells (Kang et al., 2003; Padua et al., 2008; Thiery, 2002), inducing macrophage differentiating towards a TAMs phenotype(Zhang et al., 2016a), and affecting the behavior of cancer-associated fibroblasts (Herrera et al., 2014; Stanisavljevic et al., 2015). By visualizing the view corresponding to the TGFB1:(TGFBR1+TGFBR2) pair in the multi-view CE network and calculating the degree centralities of the network, we found that there is active TGFB1:(TGFBR1+TGFBR2) communication in the leading edge between parenchyma (mainly including tumor cells) and stroma (mainly including stromal cells and immunocytes) (**Figures 2C and 2D, Methods**). We also found TGF-β1 and its receptors are widely expressed in the breast tumor tissue, and many cell types like stroma, basal-like-1, and macrophage serve as the sender and receiver of TGF-β1 signals at the same time (**Figures 2E and S2B**). This suggests that we should pay attention to how tumor cells affect other cell types such as stroma through TGF-β1 signals, not just to how other cell types affect tumor cells.

We visualized CE hotspots by eigenvector centralities and found this approach could detect core CE hotspots more efficiently (**Figure S2E**). According to the node eigenvector centrality vectors, we quantified the similarities between network views (Bródka et al., 2018) and clustered ligand–receptor pairs into four groups (**Figure S3A, Methods**). The clustering result indicates that most communications tend to be active at the leading edge between parenchyma and stroma (**Figures 2F, 2G, and S3B**).

### Decoding FCEs related to *MMP11* expression pattern

Based on the multi-view CE network constructed from the breast cancer Visium dataset, we applied HoloNet to decode FCEs. In breast cancer, cell–cell communication plays an important role in multiple biological processes such as tumor growth, invasion and migration, and leads various neoplastic and non-neoplastic cells in tumors to form complex heterogeneity (Hood and Cheresh, 2002; Lim et al., 2018). However, there still lacks the understanding of how the spatial expression patterns of specific functional genes are affected by cell–cell communication. We applied HoloNet on *MMP11* as an example (**Figure 4A**). The gene encodes matrix metallopeptidase 11 and is associated with worse overall survival through involving the degradation of the extracellular matrix (Basset et al., 1990; Zhang et al., 2016b). Previous research had found *MMP11* is expressed in stromal cells surrounding neoplastic cells (Min et al., 2013; Zhang et al., 2016b), but it is unclear how neoplastic cells trigger the spatial expression patterns of *MMP11*.

**Figure 3.**
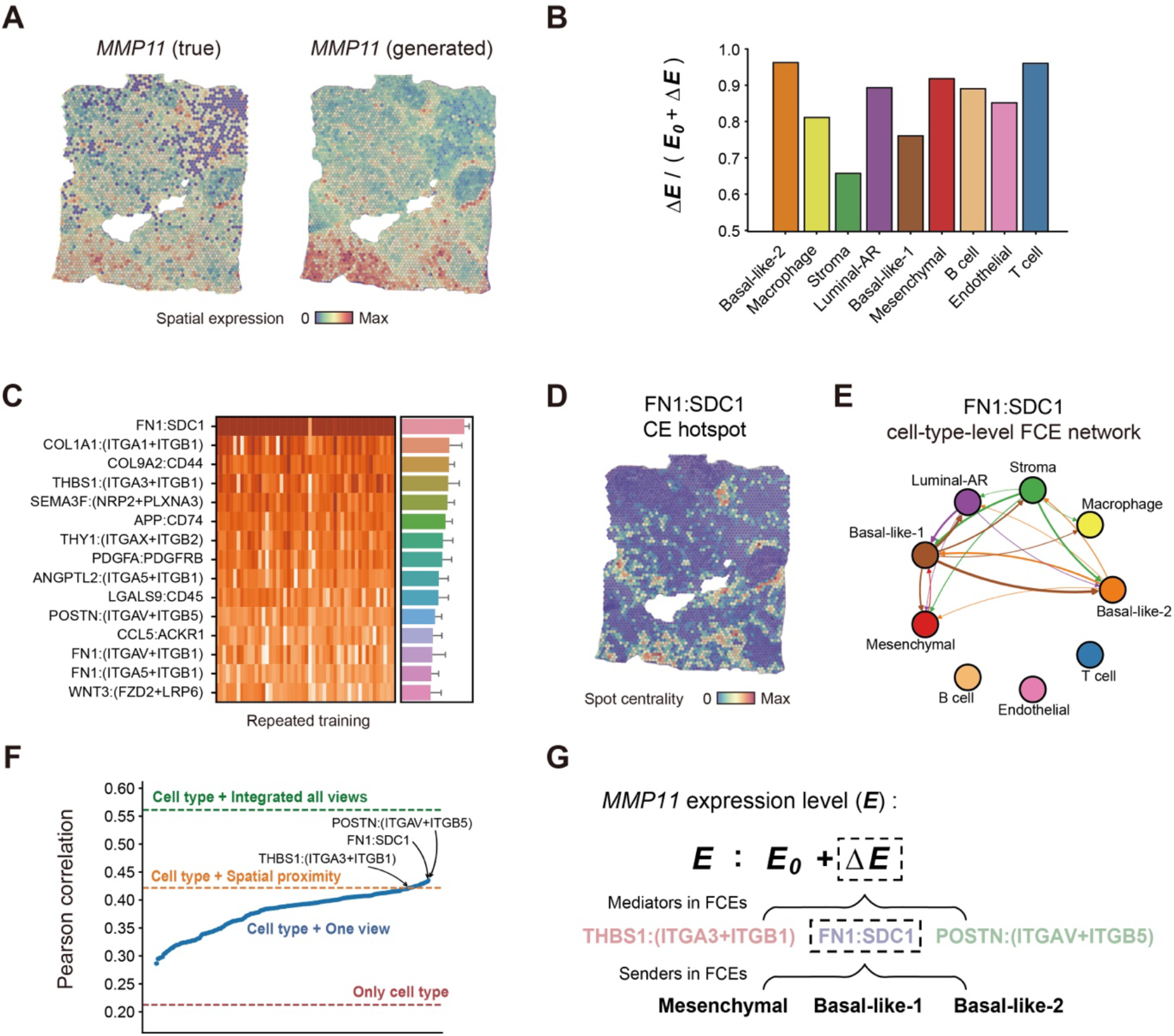
Multi-view graph learning reveals the FCE network of how cell–cell communication affects the gene expression pattern of *MMP11*. **(A)** The true (left) and HoloNet-generated (right) expression profile of *MMP11*, preprocessed by log-normalization. **(B)** The ratio of the expression change caused by CEs (**Δ*E***) to the sum of ΔE and the baseline *MMP11* expression (E_0_) in each cell type. **(C)** Top 15 ligand–receptor pairs (325 pairs in total) with the highest view attention weights in the model generating *MMP11* expression profile. The heatmap displays the attention weights of each view obtained from repeated the training procedure 50 times. The bar plot represents the mean values of the attention weights of each view, and the error bars represent standard deviations. **(D)** The CE hotspot plot of FN1: SDC1, showing the degree centrality of each spot in the FN1: SDC1 CE network. **(E)** Cell-type-level FN1:SDC1 FCE network for *MMP11*. The thickness of an edge represents the strength of *ΔE* contributed by FN1:SDC1 between the two cell types connected by the edge. **(F)** Model performances in generating *MMP11* expression using each single-view network (blue dots). We ranked the ligand–receptor pairs (x-axis) according to the model performances. The horizontal lines indicate the performances of the models only using cell-type labels (red), using cell-type labels and spatial proximity network (yellow), and using HoloNet (green). **(G)** The mechanism of multiple factors affecting *MMP11* expression revealed by HoloNet.

**Figure 4.**
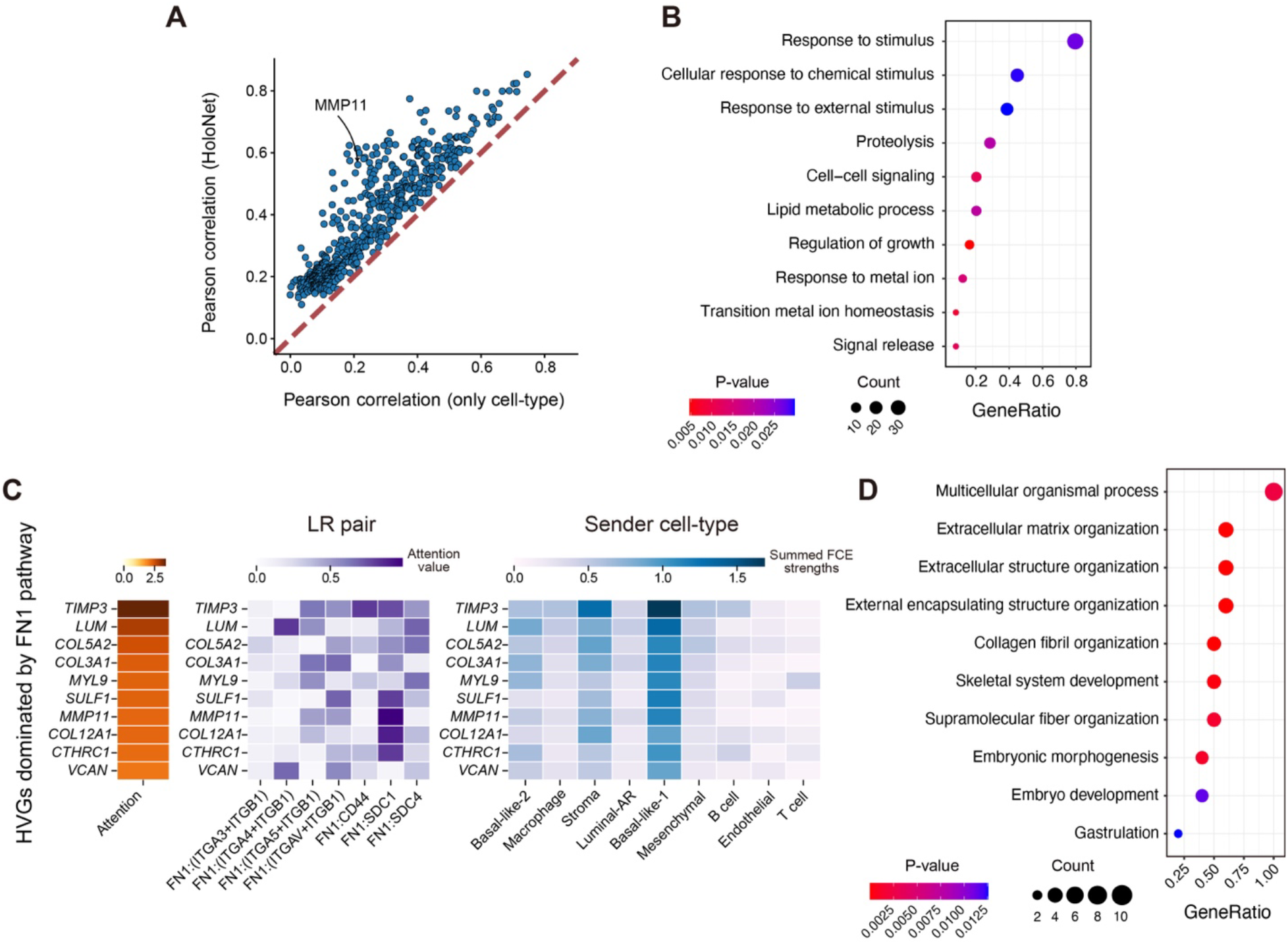
HoloNet detects genes dominated by cell–cell communication. **(A)** Performance comparison between HoloNet and models only using cell-type information. The comparison is based on tasks for generating the expression level of each gene. **(B)** Gene ontology (GO) enrichment for the top 20% genes with relatively higher performance improvements after considering CEs. **(C)** Heatmaps for FN1 pathway related genes, and which ligand receptors affect these genes, and these FCEs come from which cell types. **(D)** GO enrichment for the 10 genes related to FN1 pathway.

We trained HoloNet to generate the expression of *MMP11* and interpreted the trained model. As shown in **Figure 3A**, the expression profile of *MMP11* generated by HoloNet is quite similar to the pattern observed in the true spatial transcriptomic data. We noticed that *MMP11* is highly expressed in the lower-left part of the slide, where some stromal cells mix into tumor cells (**Figure 3A**). To identify FCEs related to *MMP11* expression, we interpreted the multi-view graph neural network of the trained HoloNet by decomposing *MMP11* expression into baseline expression *(**E_0_**)* and expression change caused by CEs (**Δ*E***). From **Figure 3B**, we can see that the expressions of *MMP11* in all cell types are mainly contributed by **Δ*E***. We found that the ligand–receptor pair FN1:SDC1 always has the highest view attention weight in the multi-view network across multiple training replicates (**Figure 3C**). This indicated that FN1:SDC1 could be one of the core ligand–receptor pairs affecting *MMP11* expression. Besides, the CE hotspots of FN1:SDC1 showed to highly coincide with the high-*MMP11* region (**Figure 3D**). Based on the cell-type-level FN1:SDC1 FCE network, we found that among all the edges linked to stroma, the one from basal-like-1 has the highest strengths, revealing that the basal-like-1 serves as the major senders of FN1 signals for triggering the change of *MMP11* expression level in stromal cells (**Figure 3E**). The spatial plots of the FN1:SDC1 FCE strengths for all cell types (**Figure S4**) show that the general FCEs from basal-like-1 cells form a spatial pattern similar to the actual expression pattern of *MMP11*. We also generated the *MMP11* expression using several single-view CE networks and discovered that the CE with FN1:SDC1 pair contributes more to determining the *MMP11* expression patterns than cell types or niches (**Figure 3F**).

The above results suggested that the FN1:SDC1 pair may regulate the expression of *MMP11*, which is consistent with previous studies. Fibronectin (FN) is a large glycoprotein as a component in the extracellular matrix and encoded by *FN1*. Fibronectin is involved in the development, invasion and migration of multiple types of human cancer (Hu et al., 2004; Jia et al., 2010; Waalkes et al., 2010; Zheng et al., 2007). Syndecans, encoded by SDC1, act as adhesion receptors play important roles in the signaling transduction (Lim et al., 2003; Woods et al., 2000; Yang and Friedl, 2016). FN has been found inducing the expression of matrix metalloproteases (MMPs) in multiple types of human cancer including the breast cancer (Das et al., 2008; Jia et al., 2004; Maity et al., 2011; Saad et al., 2002; Sen et al., 2010). In direct agreement with our results, Fernandez-Garcia et al. found that breast tumors with high FN expression tend to express higher levels of matrix metalloproteinase genes such as *MMP11*, and have a stronger metastasis tendency (Fernandez-Garcia et al., 2014). They proposed tumor-cell-derived FN can induce *MMP11* expression in mononuclear inflammatory cells (Fernandez-Garcia et al., 2014). We also detected that tumor-cell-derived FN affects *MMP11* expression in macrophages (**Figure 3E**), consistent with the previous finding. In addition to macrophages, we also found *MMP11* is mainly expressed in stromal cells and tumor cells based on the spatial transcriptomic dataset. The results from HoloNet reveals the role of various cell types in the process of FN inducing *MMP11* (**Figure 3E**), which improve our understanding of the process. We also verified that the expression level of *MMP11* has a positive correlation with the expression levels of *FN1* and *SDC1* in breast cancer samples from TCGA data (**Figures S5A and S5B; Methods**).

We also identified some other ligand–receptor pairs that have been reported to be related to *MMP11* in previous studies. For example, we found that THBS1:(ITGA3+ITGB1) and POSTN:(ITGAV+ITGB5) serve as core ligand–receptor pairs affecting *MMP11* expression (**Figures 3C and 3F**), and these signals altering *MMP11* expression in stromal cells are also mainly contributed by basal-like-1 cells (**Figures S5C-S5F**). In previous studies, THBS1 has been shown to be able to stimulate *MMP11* expression in stromal cells and enhance oral squamous cell carcinoma invasion (Pal et al., 2016). Researchers have reported that *THBS1, THBS2, POSTN, MMP11* can reflect the invasion phenotypes of breast tumors (Kim et al., 2010; Schuetz et al., 2006). All those reported results supported the *MMP11*-related FCEs identified by HoloNet. In summary, HoloNet suggested a putative mechanism of how cell–cell communication affects *MMP11* expression: basal-like-1 cells serve as a major source of *MMP11-level* alteration in stromal cells via ligand–receptor pairs such as FN1:SDC1, THBS1:(ITGA3+ITGB1) and POSTN:(ITGAV+ITGB5), which may be related to the alteration of stromal cells surrounding tumors into invasion-associated phenotypes (**Figure 3G**).

### Characterizing the FCE landscape in breast cancer

We applied HoloNet to all 586 highly variable genes (HVGs) to study how these genes are affected by cell– cell communication (**Methods**). For each target gene, we focused on the top 15 ligand–receptor pairs with the highest view attention weights and inferred the FCE network for each ligand–receptor pair (see **Data Avaliability** for details). To prove that HoloNet effectively extracts cell–cell communication information from the spatial transcriptomic data, we compared the accuracy of gene expression prediction between models only using cell-type information and those considering CEs (**Methods**). We found that the prediction accuracy on almost all genes were improved by considering CEs (**Figure 4A**). Some genes showed higher improvements, indicating that they are more affected by CEs. The top 20% genes out of 586 genes with the highest improvements are enriched in gene ontology (GO) terms such as “cell–cell signaling” and “regulation of growth” (**Figure 4B, Methods**). Similar to the case study on *MMP11*, for more than 90% of the these genes, the ligand–receptor network brought better performances than the spatial proximity network, indicating ligand–receptor networks depicted the tissue microenvironments better than spatial proximity (**Figure S6A**). Meanwhile, HoloNet achieved a better performance than the Lasso regression model using both ligand– receptor communication and cell-type information (**Figure S6B**). This results showed that the graph model of HoloNet represents the cell–cell communication mechanisms in an effective way.

We then identified the genes dominated by specific signaling pathways using HoloNet. Taking the FN1 pathway mentioned earlier as an example, FN1 pathway contains 7 ligand–receptor pairs, and our analysis revealed that some HVGs are dominated by FN1 pathway, including *MMP11* and *TIMP3* (**Figure 4C**). This result is also consistent with the findings of Fernandez Garcia et al. that FN affects the expression of MMPs and TIMPs in breast cancer (Fernandez-Garcia et al., 2014). These genes are enriched in GO terms such as “extracellular matrix organization” and “collagen fibril organization” (**Figure 4D**). We also found these genes tend to be affected by different ligand-receptor pairs, for example, *MMP11* is mainly affected by FN1:SDC1, while *TIMP3* is mainly affected by FN1:CD44 (**Figure 4C**). The FCEs for these target genes are mainly from basal-like-1 cells (**Figure 4C**). These findings revealed how FN1 pathway affects the extracellular matrix in breast tumor.

To verify the findings based on the breast cancer Visium dataset available on 10x Genomics website, we also applied HoloNet on another breast cancer Visium dataset from a publication of Wu et al. (Wu et al., 2021b). We obtained the cell-type information of this dataset in the same way as the previous dataset (**Figure S7A, Methods**), and trained HoloNet to generate the expression profiles of 285 HVGs. We generated the expression profile of *MMP11* in this dataset (**Figure S7B**). Similar to the findings on the dataset from 10x Genomics website, we also detected the layer corresponding to FN1:SDC1 with the highest attention weight, and basal-like-1 cells serve as core senders in FCEs for *MMP11* (**Figures S7C-S7F**). Similar to the results derived from the first dataset, we found that nearly all target genes are better generated when considering the CEs (**Figure S7G**), and the top 20% genes out of 285 with the highest accuracy improvements are also enriched in communication-related GO terms such as “cell population proliferation” (**Figure S7H**). These results verify the stability of the biological discovery of HoloNet between different datasets.

### Decoding FCEs in liver cancer

We applied HoloNet on a liver cancer Visium dataset (Wu et al., 2021a) (**Figure 5A**). We trained HoloNet to generate the expression profiles of 225 HVGs (**Figure 5B**). For each target gene, we focused on the top 15 ligand–receptor pairs with the highest view attention weights and inferred the FCE network for each ligand– receptor pair (see **Data Avaliability** for details). We chose *MCAM* as a case study (**Figure 5B**). *MCAM* encodes CD146, which is overexpressed in most type of tumors (Wang et al., 2020). CD146 plays an important role in cell adhesion and signal transduction processes, especially in tumor angiogenesis (Chen et al., 2017; Gao et al., 2017; Jiang et al., 2012; Tu et al., 2015; Ye et al., 2013).

**Figure 5.**
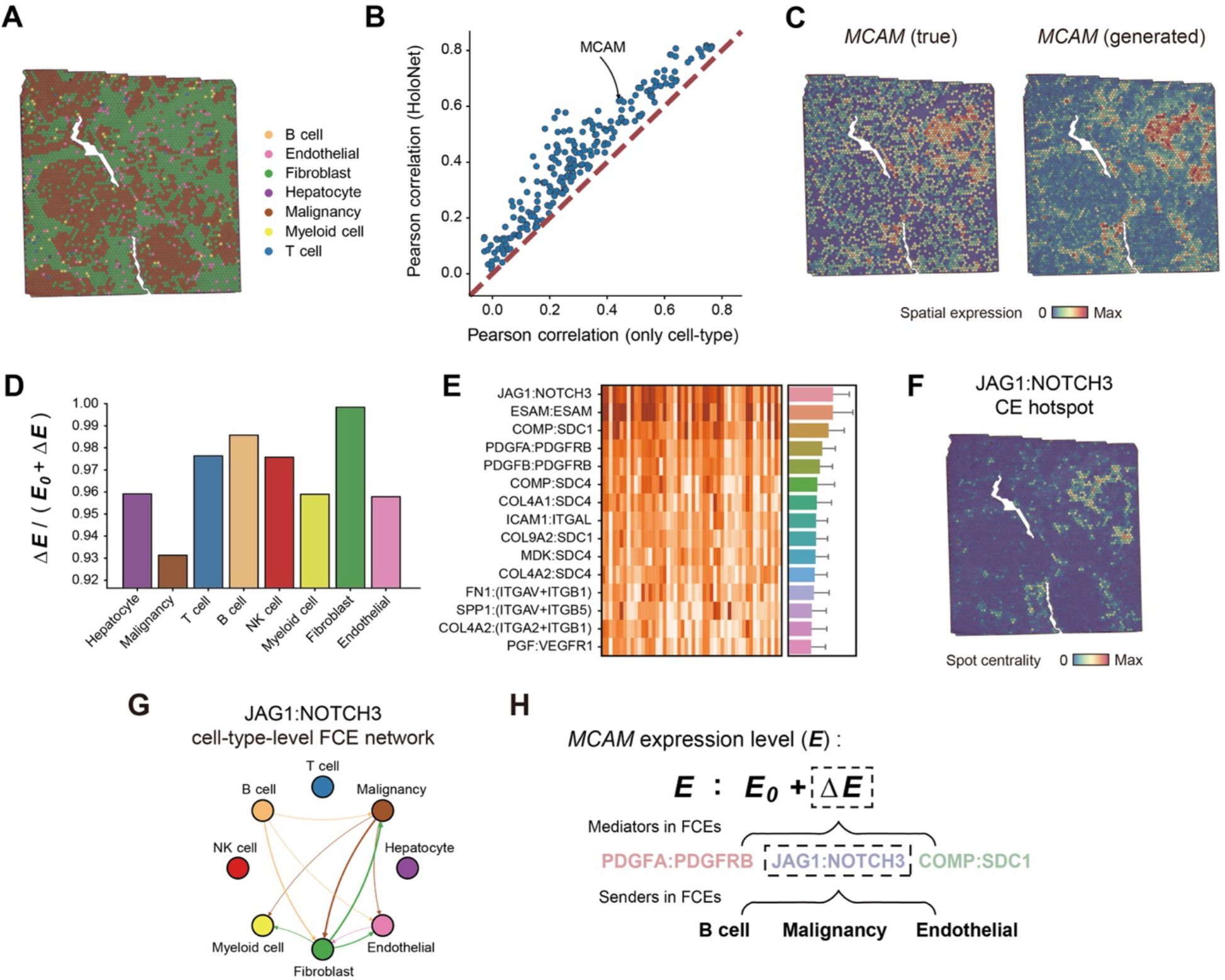
HoloNet reveals the FCE network of how cell–cell communication affects *MCAM* expression pattern. **(A)** The cell type with the highest percentage in each spot. **(B)** Performance comparison between HoloNet and models only using cell-type information. The comparison is based on tasks for generating the expression level of each gene. **(C)** The true (left) and HoloNet-generated (right) expression profiles of *MCAM*, preprocessed by log-normalization. **(D)** The ratio of the expression change caused by CEs (**Δ*E***) to the sum of **Δ*E*** and the baseline *MCAM* expression (*E*_0_) in each cell type. **(E)** Top 15 ligand– receptor pairs (202 pairs in total) with the highest view attention weights in the model generating *MCAM* expression profile. The heatmap displays the attention weights of each view obtained from repeating the training procedure 50 times. The bar plot represents the mean values of the attention weights of each view, and the error bars represent standard deviations. **(F)** The CE hotspot plot of JAG1:NOTCH3. **(G)** Cell-type-level JAG1:NOTCH3 FCE network for *MCAM*. **(H)** The mechanism of multiple factors affecting *MCAM* expression revealed by HoloNet.

We observed that *MCAM* is highly expressed in endothelial cells and fibroblasts around endothelial cells in the spatial data (**Figures 5C and S8A**). As endothelial cells cover the inner surface of blood vessels, this result indicates that fibroblasts around blood vessels have high expression of *MCAM*. To explore how cell–cell communication triggers the *MCAM* expression in stromal cells around blood vessels, we trained HoloNet to generate the expression of *MCAM* (**Figure 5C**). After decomposing *MCAM* expression into baseline expression ***E*_0_** and expression change **Δ*E*** caused by CEs, we found the expressions of *MCAM* in all cell types, especially in fibroblasts, are dominantly contributed by **Δ*E*** (**Figure 5D**). We found that JAG1:NOTCH3 and PDGFA:PDGFRB are always given the highest view attention weights (**Figures 5E and 5F**), indicating they could be the core ligand–receptor pairs affecting the expression of *MCAM*. Based on the cell-type-level JAG1:NOTCH3 FCE network, we found that the fibroblasts, malignancy cells, and endothelial cells serve as the major senders of JAG1 signals that trigger the *MCAM* expression level change in fibroblasts (**Figure 5G**). Similar results were found in the analysis of other three hepatocellular carcinoma samples from the study of Wu et al. (**Figure S9A-S9C**).

The putative mechanism of how communication affects the *MCAM* expression in fibroblasts around blood vessels is summarized in **Figure 5H**. Previous research also reported that NOTCH signals play important roles in vascular development (Dou et al., 2012; Giovannini et al., 2021; Huang et al., 2019), and JAG1 has been found involving in HCC angiogenesis as a key ligand (Nijjar et al., 2002). Pinnix et al. also found that activated Notch can directly induce *MCAM* expression in melanocytes, and multiple CSL/Notch-binding sequences in the *MCAM* promoter may mediate the process (Pinnix et al., 2009). We also verified that the expression level of*MCAM* has a positive correlation with the expression levels of *JAG1* and *NOTCH3* in liver cancer samples from TCGA data (**Figure S8B, Methods**).

We established links between pathways and all HVGs in this dataset. We identified that some HVGs, including *COL18A1* and *MCAM*, are related to the NOTCH pathway (**Figure S8C**). These genes are enriched in GO terms such as “angiogenesis” and “blood vessel development” (**Figure S8D**). In this liver cancer tissue, the functional NOTCH signals are mainly composed of JAG1:NOTCH1 and sent from fibroblasts (**Figure S8C**). These findings revealed the effect of the NOTCH pathway on angiogenesis in liver cancer.

### Validating the stability of HoloNet in a Slideseq-v2 dataset

We applied HoloNet on a mouse hippocampus Slideseq-v2 dataset (Stickels et al., 2021) to verify whether the results of HoloNet are affected by the lower resolution and sampling conditions of Visium and other spatial technologies. The dataset profiles 4,000 gene expressions of 41,786 spots with 10μm spatial resolution (**Figure 6A**). These spots with approximate single-cell resolution are assigned to 14 cell types according to their marker gene expressions (Stickels et al., 2021). We independently downsampled 5,000 spots from these spots for 3 times and applied HoloNet on the downsampled dataset. We calculated the correlation among the attention values of each ligand receptor pair from multiple down-sampling experiments, and found the attention values are stable, with a Pearson’s correlation coefficient (PCC) of 0.96, (**Figure 6B**). In a similar way, we found that the obtained FCE network also showed good stability (PCC = 0.68, **Figure 6B**).

**Figure 6.**
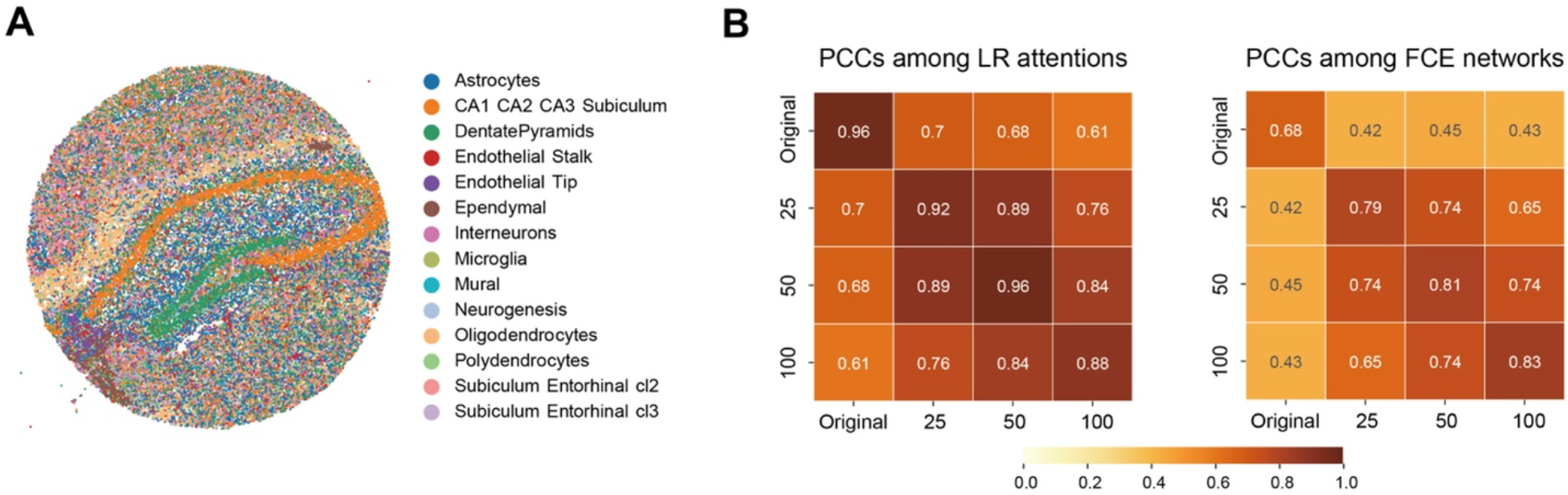
Validating the stability of HoloNet in the mouse hippocampus Slideseq-v2 dataset. **(A)** The cell type of each spot in the Slideseq-v2 dataset. **(B)** Pearson’s correlation coefficient (PCC) among LR attentions and FCE networks. ‘Original’ represents the results obtained by running HoloNet on datasets of 5000 spots after downsampling. ‘25’, ‘50’ and ‘100’ respectively represents the results obtained by running HoloNet on datasets of 5000 spots after fusing 10μm spots within 25, 50, and 100μm.

We then fused 10μm spots in certain ranges (25, 50, and 100μm) to imitate the spots in Visium datasets. We found that the attention value of each ligand-receptor pair and FCE network obtained good stability (PCC of 0.88-0.92 and 0.79-0.83, respectively, **Figure 6B**). The results obtained from different fusing ranges also have good correlation (**Figures 6B, S10A and S10B**). We noticed that the correlations between the results from fused datasets and the original dataset are not so high. This may be because the gene expression of the original data set is very sparse, resulting in a low density of the constructed communication network, which affects the performance of HoloNet (**Figure S10C**). Also, datasets with fused spots contain more comprehensive cell type composition in local tissues, which may also have impacts on the results. On the whole, HoloNet results have good stability between different sizes of spots, different sampling processes, as well as different sequencing technologies.

## Discussion

We developed a machine-learning method HoloNet to decode functional cell–cell communication events (FCEs) with spatial transcriptomic data. In HoloNet, we designed a multi-view network to model cell–cell communication events (CEs), built an attention-based graph learning model on the network to generate the target gene expression, and then interpreted the model to decode the functional CEs (FCEs). HoloNet provides the putative FCE network of how communication affects specific gene expression patterns, and reveals the four core elements of FCEs: which cell types act as senders and receivers, where they are, and which ligand– receptor pairs they employ to induce what kind of downstream responses. We applied HoloNet on two breast cancer and a liver cancer Visium spatial transcriptomic datasets to illustrate its ability to offer new biological understandings. In breast cancer, we detected ligand–receptor signals triggering the expression changes of some invasion-related genes in stromal cells surrounding tumors and identified a subtype of neoplastic cells serving as the primary senders of these signals. In liver cancer, we identified key FCEs that affect angiogenesis. Besides, we validated the stability of HoloNet using a Slideseq-v2 dataset.

Revealing the communication, cooperation, and mutual influence among various cells in tissues is important for understanding the operation of multicellular organisms and the development of diseases. We proposed the concept of FCEs as the key to answer the above questions. FCEs are the CEs that cause expression changes of specific downstream genes. Compared with previous concepts, such as ligand-target links or sender-receiver dependencies, FCE provides a basis for separating and describing the shaping of cellular phenotypes by environment from various perspectives. FCEs allows us to better characterize the whole process of intercellular communication from the sender cell to the receiver cell via the ligand–receptor pairs leading to specific downstream responses. Through the FCE-centric approach, HoloNet for the first time integrated ligand– receptor pairs, cell-type spatial distribution and downstream gene expression into a deep learning model. The model can draw holographic cell–cell communication networks on the single-cell level which could help to decode how each cell is affected by specific surrounding cells and ligand–receptor pairs.

We have developed HoloNet as a user-friendly computational tool. It is developed as a Python package that can be easily installed and customized to facilitate the systematic exploration of CEs and FCEs in user-provided datasets. Users can follow our workflow: constructing the CE network among single cells, visualizing cell–cell communications based on the network, generating the expression patterns of genes of interest, identifying the major ligand–receptor pairs and sender cell types in FCEs, and detecting the genes more affected by cell–cell communication. The main results in the article can be generated by our public python package.

HoloNet can help users solve many biological problems by revealing a FCE network that affect the expression of genes of interest. For example, similar to the analysis presented in this article, users can use HoloNet to analyze how the expression profiles of cells are affected by other cell types in the microenvironment. Provided with pre- and post-treatment spatial transcriptomic data, users can also analyze how the treatment affects the full landscape of cell–cell communication within the tissue, understand unexpected treatment outcomes, and develop new disease interventions. With the widespread use of spatial transcriptomic techniques, HoloNet can be employed in an increasing number of biological scenarios to understand the FCE network in tissues and to analyze how cellular phenotypes are affected by FCEs.

The core structure of HoloNet makes it possible to analyze various kinds of spatial transcriptomic data. In addition to 10x Visium, we applied HoloNet on another spot-based technology Slideseq-v2, and the results indicated that HoloNet has good stablility between different sizes of spots, different sampling processes, and different sequencing technologies. Single-cell-based spatial transcriptomic technologies such as SeqFISH and MERFISH can also be easily adopted as the input of HoloNet. Though, the current version of HoloNet acquires an adequate amount of cells or spots to train the GNN model, while the number of cells detected by SeqFISH (Eng et al., 2019) is too small to be well analyzed in HoloNet. Meanwhile, the number of genes profiled by MERFISH (Chen et al., 2015) is too small, so that HoloNet cannot construct CE networks using ligand and receptor gene expressions.

The current spatial technologies, especially 10x Visium, still have relatively low spatial resolution and need to use deconvolution methods to analyze cell types. HoloNet depends on the results provided by these deconvolution methods, and the results will inevitably be influenced by the deconvolution way. The simulation experiment on the Slideseq-V2 dataset showed that the the results of HoloNet using the deconvolved cell-type labels are similar with the results using the real cell-type labels. This verifies that HoloNet is not sensitive to the deconvolution method. We will focus on new spatial transcriptomic technologies with higher spatial resolution, more gene and cell numbers, and continue to update HoloNet along with the development of technologies.

In the future, we can incorporate prior knowledge embedding into the model, such as the intracellular signaling networks or known biological functions of some ligands, to better reveal how FCEs affect downstream gene expressions. In addition, with the development of spatial sequencing technologies, more subdivided cell types or positions on pseudotime trajectories can be used here to implement multi-scale models of FCEs and gain deeper biological understandings.

## Methods

### Datasets and data pre-processing

We applied HoloNet on four datasets, including two human breast cancer Visium spatial transcriptomic datasets, a human liver cancer Visium spatial transcriptomic dataset and a mouse hippocampus Slideseq-v2 dataset. The main breast cancer dataset in this article is from the 10x Genomics website (https://www.10xgenomics.com/resources/datasets, Visium Demonstration, Human Breast Cancer, Block A Section 1) and profiled the expression of 24,923 genes in 3,798 spots on a fresh frozen invasive ductal carcinoma breast tissue sample. The breast cancer dataset for validation is obtained from the publication of Wu et al.(Wu et al., 2021b). They performed spatial transcriptomics on six samples, and we selected the 1160920F sample considering the cell-type richness and the number of spots. The dataset includes the expression profiles of 28,402 genes in 4,895 spots. We excluded spots with less than 500 UMIs and genes expressed in less than 3 spots for both datasets and then normalized the expression matrix with the *LogNormalize* method in Seurat(Stuart et al., 2019). We annotated the cell types by label transfer (the *TransferData* function in Seurat) using a well-annotated single-cell breast cancer dataset (accession code is GSE118390) (Karaayvaz et al., 2018) as reference. The reference dataset contains four cell types of malignant components (basal-like-1, basal-like-2, luminal-AR, and mesenchymal) and five cell types of stromal and immune components (macrophage, B cell, T cell, stromal, and endothelial cell). The liver cancer dataset is s obtained from the publication of Wu et al. (Wu et al., 2021a). We selected the “HCC_2T” sample from tumor tissue samples as this sample has a high number of spots. The dataset profiled the expression of 33,538 genes in 4,733 spots. We also used the “HCC_1T”, “HCC_2T” and “HCC_4T” samples for validation. Referring to the signature-based cell type scoring strategy of Wu et al., we used their curated gene signatures of common cell types in liver cancer and defined the average log-normalized expression values of the genes in the signature as the corresponding cell type scores. As for the Slideseq-v2 dataset, the mouse hippocampus Slideseq-v2 dataset includes the expression profiles of 4,000 genes in 41,786 spots with a spatial resolution of 10 μm (Stickels et al., 2021). The Squidpy package provided the preprocessed mouse hippocampus Slideseq-v2 dataset with celltype labels (Palla et al., 2022), and we run HoloNet based on the dataset.

### Multi-view CE network

Inspired by the concept of multi-view social networks (Bródka et al., 2018; Kivelä et al., 2014; Shi et al., 2016), we used a node-aligned directed weighted multi-view network to represent CEs in spatial transcriptomics. Formally, a multi-view CE network *G* with a set *V* of views is defined on the aligned node set *U*. The node set *U* represents single cells (or spots) in the measured tissue. Each view *v* ∈ *V* is a directed graph *G^(v)^* = (*U, E^v^*), in which *E^(v)^* consists of edges between nodes, and each edge 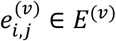 has a weight 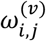 for *i, j* ∈ *U*. The edge weight 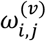 is calculated by a designed edge weighting strategy, representing the strength of CEs from cell *i* to cell *j* mediated by a specific ligand–receptor pair corresponding to the view *v*. The multi-view CE network can be represented by a 3-dimension CE tensor (cell by cell by ligand–receptor pair) as a set of adjacency matrices *A^(v)^*. The edge weight 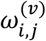 is the (*i, j, v*) element in the CE tensor. To optimize the computational efficiency, we used Pytorch(Paszke et al., 2019) to implement the multi-view network in the following steps.

### Edge weighting strategy

To calculate the edge weight 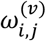, we designed a new edge weighting strategy according to the law of molecular diffusion(Balluffi et al., 2005) and the law of mass action in chemistry. According to Fick’s second law, considering a one-dimensional diffusion, the ligand concentration at *x* position and in time *t* is:

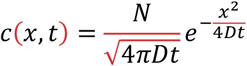

where *N* is the number of “source” atoms per unit area initially placed at *x* = 0, and *D* is the coefficient of diffusion. This inspired us to model cell–cell communication in the spatial transcriptomic data using Gaussian functions. In spatial data, considering the ligand molecules (encoded by gene *L_v_*) released by cell *i* (sender cell), the concentration of these ligand molecules at the position of cell *j* (receiver cell) is determined by the ligand expression in cell *i* and the distance parameter of cell *i* and *j*:

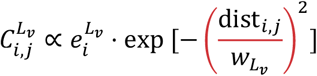

where cell *i* and cell *j* can be any cells in the spatial data, 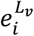 is the expression level of the ligand gene (*L_v_*) in the sender cell *i*, *dist_ij_* is the Euclidean distance between the spatial coordinates of the sender cell *i* and the receiver cell *j*, and *w_L_v__* describes the diffusion ability of the ligand encoded by *L_v_* which controls the covering region of ligands from one sender cell.

Here we set the proportionality coefficient as 1. Thus, for sender cell *i*, receiver cell *j*, and ligand–receptor pair *v*, the edge weight 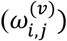 is calculated as:

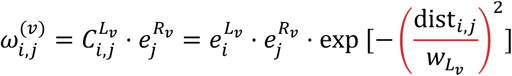

where 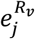 is the expression level of the receptor gene (*R_v_*) in the receiver cell *j*. When a ligand or receptor molecule is a multiple complex and controlled by multiple genes, 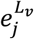 and 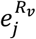 are the geometric means of these genes.

We also built up a relationship between our edge weighting strategy and the traditional strategy which only considers the cell–cell communication between neighboring cells within a fixed region. We selected *w_L_k__* to make the amount of ligand molecules diffusing throughout the whole tissue equivalent to covering a region with a diameter (*d*) at a fixed concentration (*d* = 255 μm as the default setting in Visium datasets, where the sender spot diameter is 55 μm and ligands diffuse 100 μm)(Fischer et al., 2021; Li et al., 2020). Specifically, we let

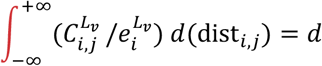

Considering

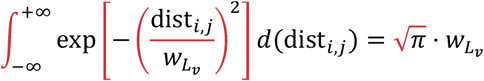

we can get

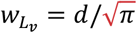

The specific value of *w_L_v__* can be selected visibly by the *select_w* function in HoloNet (**Figure S1A**). We filtered out ligand–receptor pairs in which either the ligand or the receptor expressed in less than 30% of the cells.

HoloNet provides a method to include different properties of different ligand–receptor pairs into the calculation of *w_L_v__*. We obtained the information on ligands and receptors from CellChatDB (Jin et al., 2021). The database categorizes ligand–receptor pairs into three categories: ‘ECM-receptor’, ‘secreted signaling’, and ‘cell-cell contact’. We calculated the *w_L_v__* of ‘ECM-receptor’ and ‘cell-cell contact’ ligand–receptor pairs by the aforementioned method, and restricted the communication between adjacent spots. Correspondingly, we set the *w_L_v__* of ‘secreted signaling’ ligand–receptor pairs twice as large as *w_L_v__* of ‘ECM-receptor’ and ‘cell-cell contact’ ligand–receptor pairs by default.

HoloNet also considered other important signaling cofactors. CellChatDB provides soluble agonists *(AG_v_*), antagonists *(AN_v_*), as well as stimulatory and inhibitory membrane-bound co-receptors *(RA_v_* and *RI_v_*) for each ligand–receptor pair (Jin et al., 2021). We calculate the molecular concentration of agonists *(AG_v_*) at receiver cell *j* as:

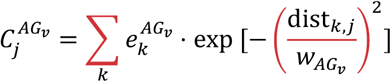

where *k* is the source cell of *AG_v_*, and 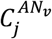 can be calculated in the same way. Then, inspired by CellChat (Jin et al., 2021), for sender cell *i*, receiver cell *j*, and ligand–receptor pair *v*, the edge weight 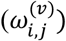 including other signaling cofactors is calculated as:

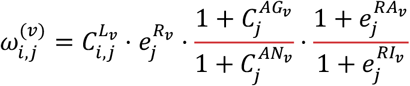

### Filtering out edges with low specificities within a multi-view CE network

Both the effects of non-specifically widely expressed ligands (or receptors) and sequencing technology errors could introduce unspecific edges into the multi-view CE network. To identify which cell pairs are actively communicating, we proposed a strategy to calculate the edge specificity within each CE network view. Developed from the stLearn permutation test strategy (Pham et al., 2020), we selected *n* background gene pairs (*n* = 200 by default) for each ligand–receptor pair. The two genes of each background gene pair have similar average expression levels to the ligand and receptor, respectively. Then we used the background gene pairs to generate the null distribution of each edge and filtered out edges whose weights are not significantly larger than the null distribution (**Figure S1B**).

Specifically, 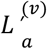 and 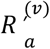 is the *a*-th pair of background genes for ligand–receptor pair *v*, which respectively exhibit close average expression levels with *L_v_* and *R_v_* in the dataset. We calculated the background edge weight between cell *i* and cell *j* based on the background gene pair:

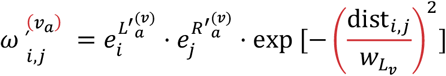

The edge weights of *n* background gene pairs 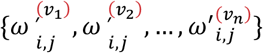 form the null distribution of 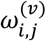. We would filter out the edge if 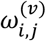 isn’t larger than enough background edge weights (threshold is set as 0.95 by default). We regarded the proportion of background edge weights that are lower than 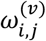 as the specificity of the edges. In the first breast cancer dataset, removing edges with low specificities reduced the network density in each view of the network, and the mean of connectivity is reduced from 0.173±0.116 to 0.046±0.050 (0.116 and 0.050 are standard deviations, **Figure S2C**). Taking TGFB1:(TGFBR1+TGFBR2) pair as an example, the detected CE hotspots cover a diffuse region before filtering as *TGFB1, TGFBR1* and *TGFBR2* are wildly expressed. The hotspots are more clearly defined at the protruding region between parenchyma and stroma after filtering out low-specificity edges (**Figures S2D and S2E**).

### Visualizing CEs based on the multi-view network

#### CE hotspot

We introduced centrality metrics from social network analysis to visualize the spots with high activities of specific ligand–receptor communication. For the view *G^(v)^* in the network *G*, which represents the CEs via ligand–receptor pair *v*, we calculated the degree centrality *dc_i,v_* and the eigenvector centrality *ec_i,v_* for the cell (or spot) *i*. We used the CE hotspot plot to visualize the centralities of spots.

We obtained the degree centrality *dc_i,v_* for cell *i* and ligand–receptor pair *v* by summing the indegrees and outdegrees of cell *i* in *G^(v)^*:

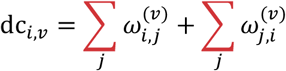

where 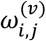 is the edge weight from cell *i* to cell *j* in *G^(v)^*. The indegrees and outdegrees reflect the receiving and sending activities respectively for the corresponding cells. We regarded the in-degree minus out-degree as the sender-receiver attributes, and plotted the attributes in **Figure 2C**.

As for the eigenvector centrality, we denoted *ec_i,v_* as 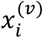 and calculated *ec_i,v_* by solving the equation:

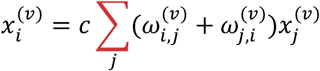

where *c* is a constant of proportionality and is set as 1 by default. To obtain a stable ***x(v)***, where 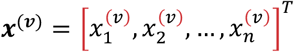, each iteration updates *x^(v)^*(*t*) according to:

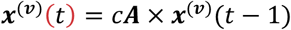

until *x^(k)^*(*t*) = *x^(k)^*(*t* – 1).

#### Cell-type-level CE network

We calculated the cell-type-level CE network to represent the general strengths of CEs from one cell type to another cell type. For ligand–receptor pair *v*, the edge weight 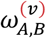 between sender cell type *A* and receiver cell type *B* is calculated by summing up the edge weights between cells belonging to cell types *A, B* in the CE network view *G^(v)^*:

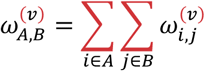

where 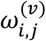 is the edge weight from cell *i* to cell *j* in *G^(v)^*.

### Ligand–receptor pair clustering

Based on the multi-view CE network, we calculated the dissimilarities between pairwise ligand–receptor pairs and identified the ligand–receptor pairs with similar spatial active regions. For two ligand–receptor pairs *u* and *v*, we calculated the dissimilarities between them:

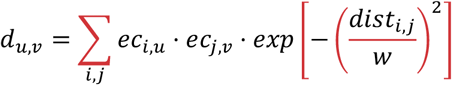

where *ec_i,u_* is the eigenvector centrality for cell *i* and ligand–receptor pair *u, dist_i,j_* is the Euclidean distance between the spatial coordinates of the sender cell *i* and the receiver cell *j*, and *w* is a factor to control the influence of distance. We set *w* same as *w_L_v__*. We used the hierarchical clustering method in sklearn package (Pedregosa et al., 2011) to cluster ligand–receptor pairs according to the dissimilarities. Then, we summed up the CE hotspots of ligand–receptor pairs in the same cluster, and obtained the cumulated spatial centrality of LR pair clusters.

### Generating specific gene expressions via multi-view graph learning

We proposed a graph neural network on the multi-view CE network *G* to generate the expression profile of specific target genes and interpreted the generating process to reveal how CEs affect these genes. One of the inputs of the model is the cell-type matrix ***X*** ∈ *R^N×C^*, where *N* is the number of cells, *C* is the number of cell types and ***X*** is the cell-type percentage of each spot or one-hot encoded cell-type labels. Besides, the multiview CE network *G* adjacency matrices *_A_* = {***A*^(1)^, *A*^(2)^,…, *A*^(*V*)^**} is another input, where ***A^(v)^*** ∈ *R^N×N^*, *N* is the number of cells and *V* is the number of views (same as the number of ligand–receptor pairs). The output of the model is the generated gene expression profile 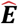. For a target gene t, the model can be described as:

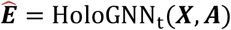

#### Selecting target genes to be generated

We selected genes to be generated by our model according to the following rules: (1) the genes should be HVGs (identified by the *highly_variable_genes* function in Scanpy package (Wolf et al., 2018) using parameters min_mean = 0.05 and min_disp = 0.5); (2) the genes should be expressed in more than 50% cells; (3) the genes should not be mitochondrion genes, ligand or receptor genes of the expressed ligand–receptor pairs. We scaled the expression levels of these genes to 0~1. When we had to use a receptor (or ligand) genes as the target gene, we removed ligand–receptor pairs containing the receptor gene and its corresponding ligand gene from the ligand–receptor pair list.

#### Preprocessing adjacency matrices

Cells of the same type that have a close spatial location may have unexpected similarities, for example they may originate from the same progenitor cell. This similarity is dependent on spatial location, but not affected by cell–cell communication, and can interfere with model results as confounding factors. We removed the edges between cells of the same type to remove the confounding factors. We calculated the cell-type similarity (*S_i,j_*) between cell *i,j* and the adjusted adjacency matrix 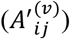 as:

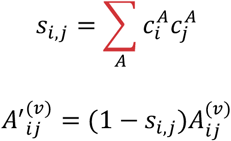

where 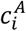 represents whether cell *i* belongs to cell type *A*, or in spot-based datasets, represents the proportion of cell type *A* in spot *i*.

Next, we calculated the normalized adjacency matrix *Â^(v)^* for each network view as:

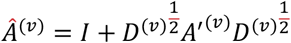

where 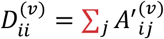

#### Graph model architecture

With the cell-type matrix ***X*** and preprocessed adjacency matrices 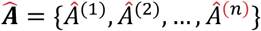 as inputs, graph models were constructed to generate the baseline expression ***E*_0_** determined by its cell type and the expression change **Δ*E*** caused by CEs, and then these two components were summed up to get the expression profile prediction 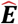 (**Figure 1C**).

To estimate **Δ*E***, we used each matrix in 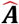 as the adjacency matrix and ***X*** as the feature matrix to construct a GNN for each view, and then obtained the embeddings of nodes in each view. Then the embeddings of nodes from each view were integrated with the attention mechanism (learnable weights *c_v_* for each view) to generate the final embeddings **Z_Δ_**:

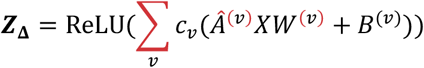

where *Â^(v)^X* ∈ *R^N×C^* (*N* is the number of cells and *C* is the number of cell types) represents the general CE strengths from each sender cell type to each cell via the ligand–receptor pair *v, W^(v)^* ∈ *R^C×H^* (*H* is the hidden layer dimension and is set as *C* by default) incorporates the effects from each sender cell type, *B^(v)^* ∈ *R^N×H^* serves as the bias matrix, *c_v_* incorporates effects from each ligand–receptor pair, and *ReLU* is an activation function:

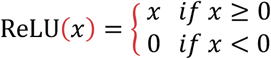

Next, we used a fully connected layer to estimate **Δ*E*** (**ΔE** ∈ *R^N^*):

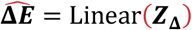

Besides, we constructed another fully connected layer to estimate ***E*_0_** by the cell-type matrix *X*:

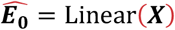

And the expression profile prediction 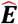 is calculated as:

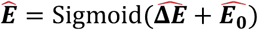

where *Sigmoid* is an activation function:

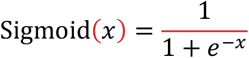

We used the mean squared error (MSE) between 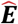 and true expression profile *E* as the loss function:

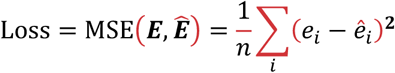

#### Training strategy

We split cells (or spots) in any dataset into train and validation sets. By default, we used 85% cells as training sets and 15% as validation sets. We trained the model for 500 epochs in the training sets and select the model with the lowest MSE in the validation sets as the final model for the following interpretation. We optimized parameters using the Adam optimizer (initial learning rate is 0.1, and weight decay is 5 × 10^-4^) and the StepLR learning rate decay strategy (step size is 10, and gamma is 0.9) by default.

To verify the robustness of the model and improve the reliability of interpretation, we repeatedly trained our graph model for certain times (50 times by default when decoding FCEs and 5 times by default when identifying target genes more affected by cell–cell communication). The final expression profile predictions displayed in the article are the mean value of expression profiles derived from each repetition.

#### Model interpretation

We obtained a trained model *HoloGNN_t_* for the target gene *t* after training. We interpreted the model in three ways:

1. **Detecting cell types more affected by FCEs.** We calculated the mean value of 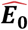 and 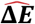 from each trained *HoloGNN_t_*. And then we calculated 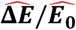 for each cell type and regarded cell types with higher 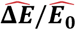 as cell types more affected by FCEs.
2. **Identifying ligand–receptor pairs mediating FCEs.** For each trained *HoloGNN_t_*, we extracted the view attention *c_v_* which represents the contribution of each view *v* (corresponding to the ligand–receptor pair *v*) to the expression profile prediction (**Figure S1C**). We calculated the mean absolute value of *c_v_* in each repeatedly trained *HoloGNN_t_* and plotted these results in the FCE mediator plot.
3. **Identifying the major sender and receiver cell types in FCEs.** As mentioned before, *Â^(v)^X* matrix represents the general CE strengths from each cell type to each cell, and *W^(v)^* incorporates the impacts from each sender cell type. Thus, we calculated the general FCE strengths from sender cell type *C* to the cell *i* as:

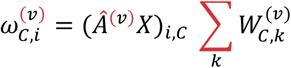 The general FCE strengths for each cell type are plotted in **Figure S3**. The computing process is also shown in **Figure S1C**. Also, we summed up the contribution of sender cell type *C* to all cells of cell type *C’*, and plotted it as the cell-type-level FCE network. The weight of the edge from cell type *C* to *C’* in the cell-type-level FCE network is calculated as:

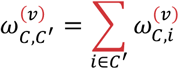

#### Generating expression profile with each network view

To verify some ligand–receptor pairs indeed facilitate cells exhibit specific gene expression patterns and remove the multicollinear effect, we generated the target gene expression profiles with each single view of the CE network. Similar to the *HoloGNN_t_*, the architecture of the model using the network view *v* is:

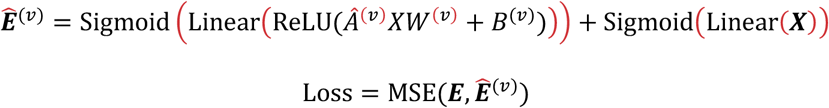

#### Baseline model

In this article, we used three baseline models: a model only using cell-type information (**Figure 4**), a model using the spatial proximity relationship matrix rather than a multi-view CE network as the adjacency matrix (**Figures 3F and S6B**), and a Lasso model with both ligand–receptor communication and cell-type information (**Figure S6C**).

As the first baseline model, we trained a full-connected network using cell type matrix *X* only, and obtain 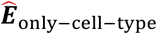 according to:

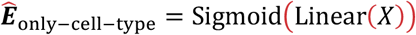

As the second baseline model, we trained a graph model similar to HoloGNN_t_ but used the spatial proximity relationship matrix as the adjacency matrix:

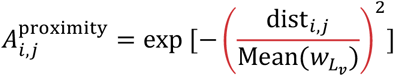

We got the preprocessed adjacency matrix *Â*^proximity^ in the way discussed before. Then we constructed a graph model and obtained 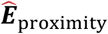 according to as:

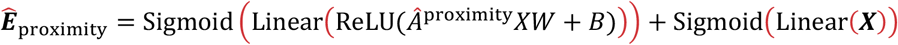

The loss function, training strategy and hyperparameters are same as the HoloNet GNN model.

As the third baseline model, we generated the target gene expression with Lasso regression (‘sklearn’ LassoCV function with default parameters) using both the ligand–receptor communication and cell-type information:

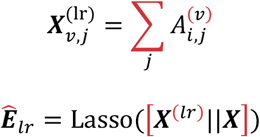

where ***X*** is the cell-type information, and 11 represents the concatenation of matrices. We calculated the Pearson correlation of 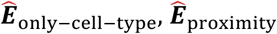 and 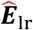 with ***E*** as the accuracies of baseline models and compared them with the accuracy of HoloNet GNN model.

#### Identifying the genes dominated by a certain signaling pathway

CellchatDB provides the corresponding signaling pathway of each ligand receptor pair. We repeatedly trained the graph model of HoloNet for 50 times for each target genes, and for a target gene *i*, we obtained the attention value 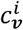 corresponding to each ligand–receptor pair (*v*). For the signaling pathway (*P*) of interest, CellchatDB provides the set of ligand–receptor pairs (*v*_1_, *v*_2_,…, *v_k_* ∈ *P*) belonging to the pathway. We obtained 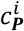 by adding up the attention values from 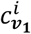 to 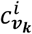, and ranked the results 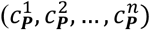 from different target genes (1,2,…, *n*). Ten target genes with the highest 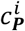 are regarded as the genes dominated by signaling pathway *P*.

In **Figure 4C** and **S8C**, the left columns show ranked 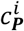 and the medium columns show 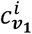 to 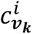. As for the right columns, we calculated the weight of the edge from cell type *C* to all cell types in the cell-type-level FCE network as 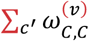, and plotted the weight in the right columns.

### Validating the stability in Slideseq-v2 datasets

The mouse hippocampus Slideseq-v2 dataset includes 41,786 spots. First, we sampled 5000 spots from them, and ran HoloNet in the dataset after downsampling. For each target gene (*i*) and ligand–receptor pair *(v)*, we obtained the attention value 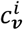 and the vectorized cell-type-level FCE network 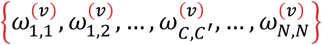, where *C* and *C*’ can be any cell type. Second, in the same way, we repeatedly sampled 5000 spots and obtained the new attention value 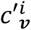 and the new vectorized cell-type-level FCE network 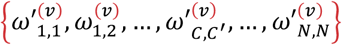. For attention values, we calculated the Pearson’s correlation coefficient (PCC) 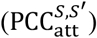 between the two experiments (named as *S* and *S*’) as:

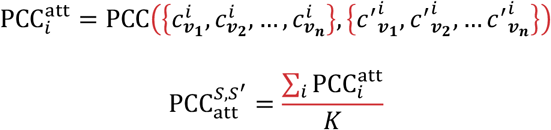

where *K* is the number of target genes. On the other hand, we calculated 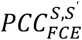 as:

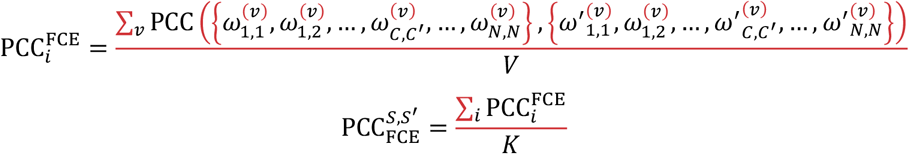

where *n* is the number of ligand–receptor pairs. We repeatedly sampled for three times (*S, S*′ and *S*″) and calculated the mean value of 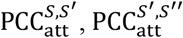 and 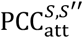 (**Figure 6B**). Also, we calculated the set of mean values of 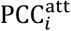 among three experiments, representing the stability of HoloNet on the task of generating different target genes (“Original vs Original” column in **Figure S9A**). We also analyzed PCCs for FCE network in the same way.

Next, we sampled 5000 spots, then regarded these spots as center spots. For each center spot, we added the gene expression values of all spots within 25μm around the center spot to the center spot. In this way, we down-sampled the dataset and fused 10μm spots within 25μm. We also repeated this process for three times. In a similar way like before, we calculated the PCC of these three repeats and between the “Original” and fused dataset. Similarly, we fused spots within 50, and 100μm and validated the stability of results.

### Gene ontology enrichment analysis for genes linked to CEs

We characterized the genes more affected or less by cell–cell communication using the ‘clusterProfiler’ R package (version 3.14.3) (Yu et al., 2012). We ranked the target genes with the performance improvements after considering CEs. We enriched the top or bottom genes into ‘biological processes’ GO terms, regarding all target genes used to be generated as the background gene set. The results of enrichment analysis are visualized with the dot plot.

### TCGA data analysis

We applied GEPIA (Tang et al., 2017) to study the gene expression data from the TCGA database. We performed the Pearson’s method to calculate correlation coefficients between expression levels of different genes in TCGA breast cancer data.

## Data availability

The used two breast cancer datasets can be downloaded from the 10x Genomics website (https://www.10xgenomics.com/resources/datasets) and Zenodo data repository (https://zenodo.org/record/4739739). The liver cancer dataset can be downloaded from https://ngdc.cncb.ac.cn/gsa-human/browse/HRA000437. The preprocessed public spatial data is available at https://zenodo.org/record/6602473. The results derived from the breast and liver cancer spatial datasets are provided on https://zenodo.org/record/7611600.

## Code availability

HoloNet is implemented as a Python package. The code and tutorial of HoloNet are available at https://github.com/lhc17/HoloNet.

## Acknowledgments

We thank Sijie Chen, Haoxiang Gao, Jiaqi Li, Chen Li, Xi Xi, Haiyang Bian, and Yixin Chen for helpful discussions.

The work is supported in part by National Key R&D Program of China grant 2021YFF1200900, NSFC grants (62050178, 61721003, 62103227), and the CZI HCA Seed Network grant 2019-02444.

## Author contributions

H.L., M.H., T.M., L.W., X.Z. conceived the study. L.W. and X.Z. supervised the research. H.L. designed and implemented the algorithm. H.L. and T.M. prepared the figures. H.L., L.W., and X.Z. wrote the manuscript.

