## Supplementary figures for "Decoding functional cell–cell communication events by multi-view graph learning on spatial transcriptomics"

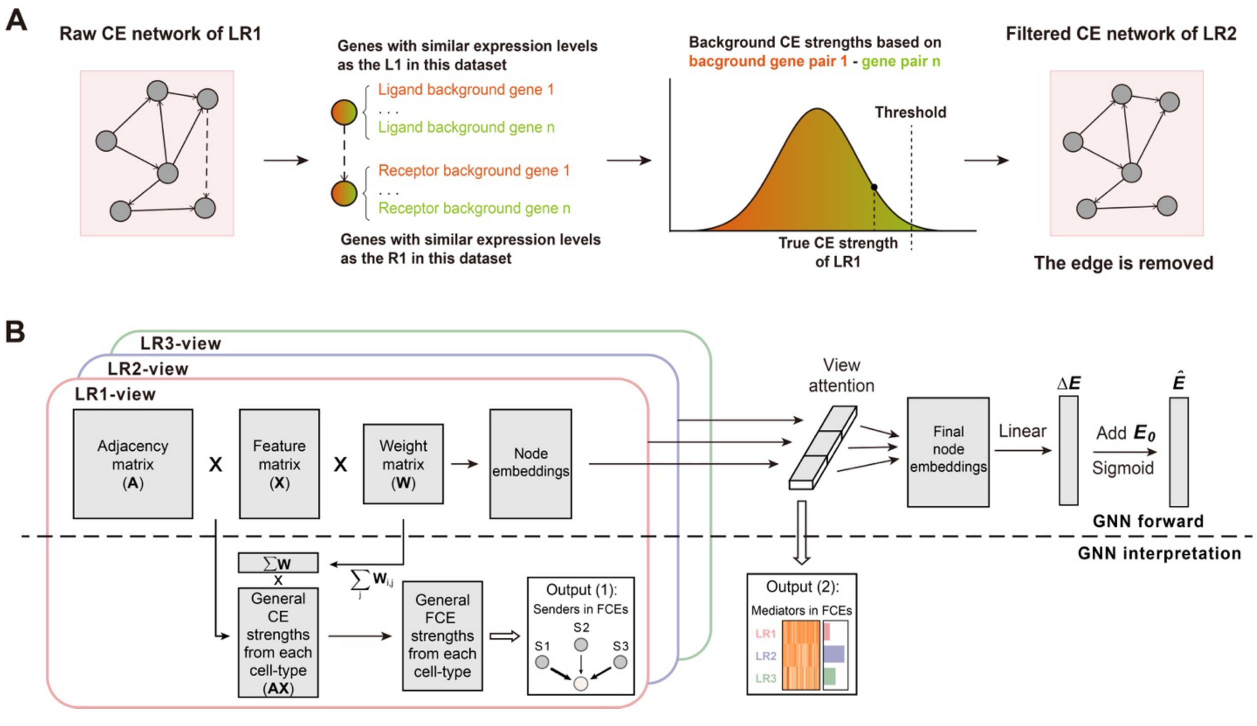


**Figure S1. Constructing a multi-view CE network.**

**(A)** Filtering false positive edges. **(B)** The detailed structure and interpretation strategy of the HoloNet GNN in a matrix form.

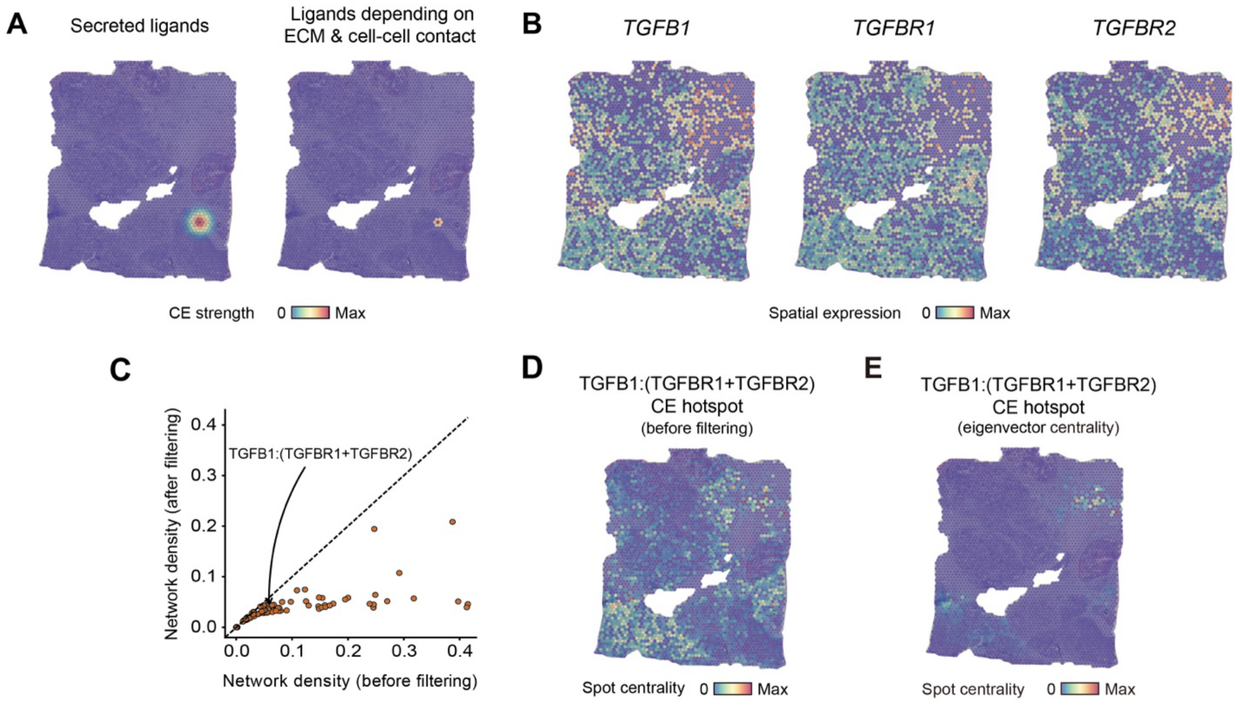


**Figure S2. Constructing and visualizing the COL1A1:DDR1 CE network.**

**(A)** Covering region of ligands from a spot. **(B)** Expression profile of *TGFB1*, *TGFB1*, and *TGFB2*. **(C)** The changes of network density before and after filtering edges with low specificities. One point indicates the change of a single-layer network based on a ligand-receptor pair. **(D)** CE hotspot plot describes the regions with active TGFB1:(TGFBR1+TGFBR2) signals, using the CE network without filtering, comparing with **Figure 2D**. **(E)** The eigenvector centrality of each spot in the CE network based on TGFB1:(TGFBR1+TGFBR2). Eigenvector centrality can better detect CE hotspots with a clear center than degree centrality in **Figure 2D**.

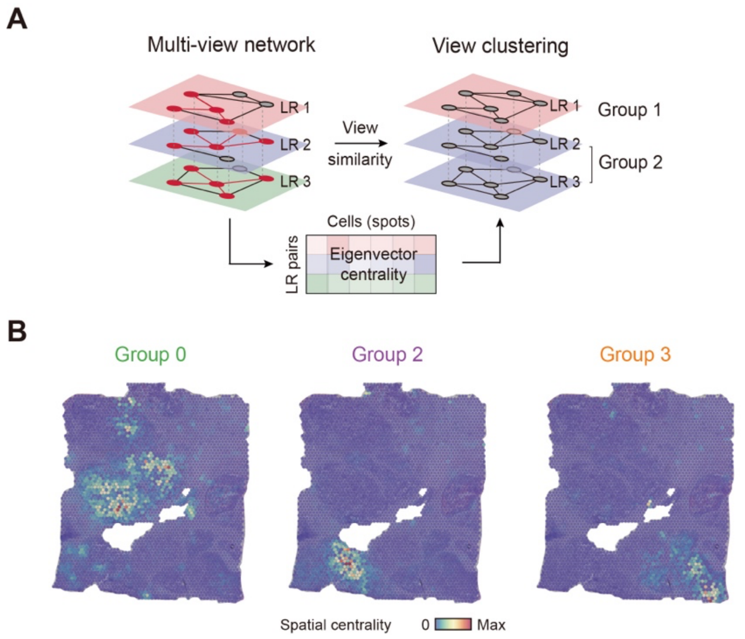


,

**Figure S3. Ligand–receptor pair clustering based on the multi-view CE network.**

**(A)** Clustering views according to the view property matrix. The view property matrix contains node eigenvector centralities (columns) of each view (rows). **(B)** General CE hotspot of each ligand-receptor group. All hotspots of members in a group are superimposed to generate the general hotspot.

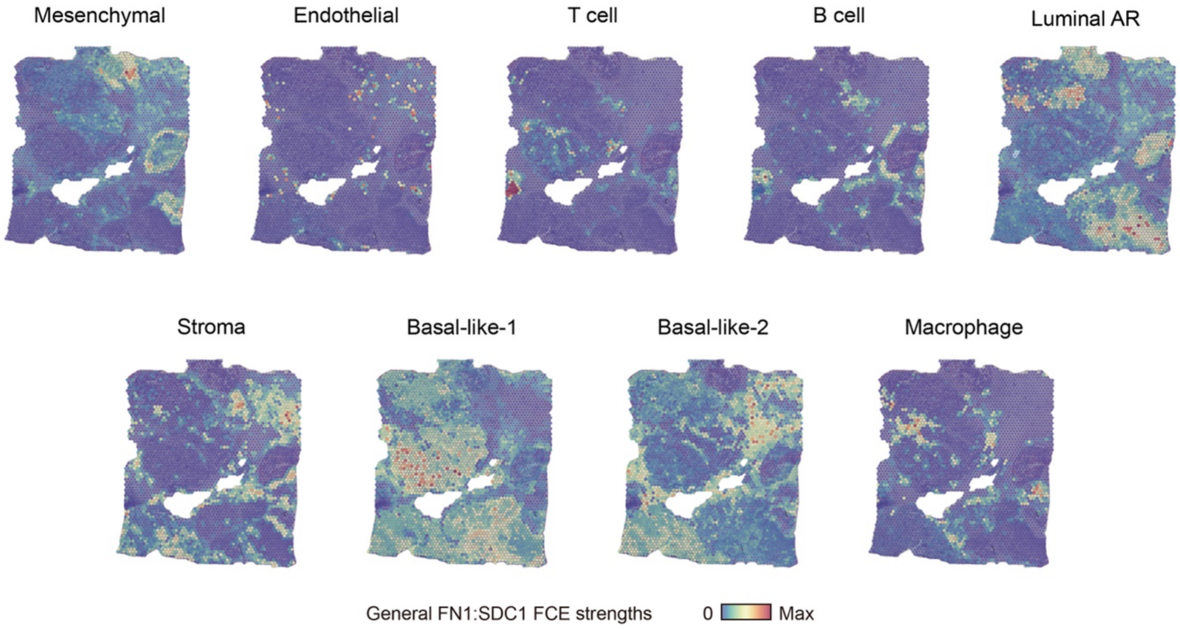


**Figure S4. The general FN1:SDC1 FCE strengths from each cell-type to each spot.** The FCE strengths in each subplot is scaled to 0-1.

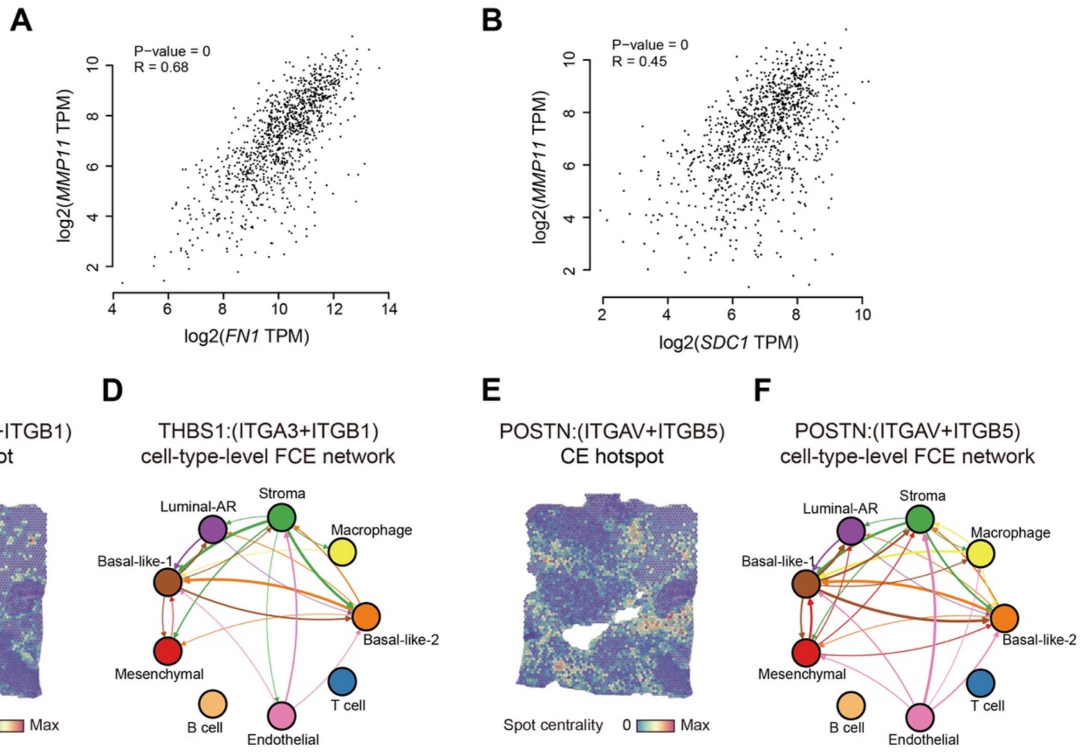


**Figure S5. FCEs related to *MMP11* expression pattern.**

**(A)** Correlation analysis of *FN1* and *MMP11* expression based on TCGA breast tumor samples. **(B)** Correlation analysis of *SDC1* and *MMP11* expression based on TCGA breast tumor samples. **(C)** The CE hotspot plot of THBS1:(ITGA3+ITGB1). **(D)** Cell-type-level THBS1:(ITGA3+ITGB1) FCE network for *MMP11*, similar as **Figure 3E**. **(E)** The CE hotspot plot of POSTN:(ITGAV+ITGB5). **(F)** Cell-type-level POSTN:(ITGAV+ITGB5) FCE network for *MMP11*, similar as **Figure 3E**.

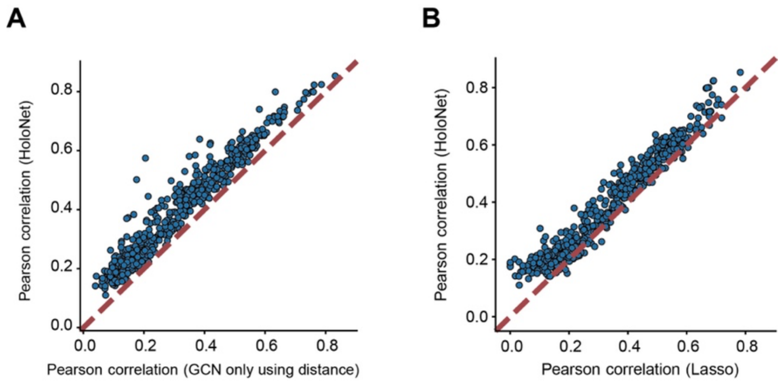


**Figure S6. The performance of HoloNet on multiple target genes.**

**(A)** Performance comparison between HoloNet and the graph model using spatial proximity network as the adjacency matrix. The comparison is based on tasks for predicting the expression level of each gene. **(B)** Performance comparison between HoloNet and Lasso regression models. The Lasso regression models predict the target gene expressions using cell-type information and the various ligand-receptor CE strength received by each cell. The comparison is based on tasks for predicting the expression level of each gene.

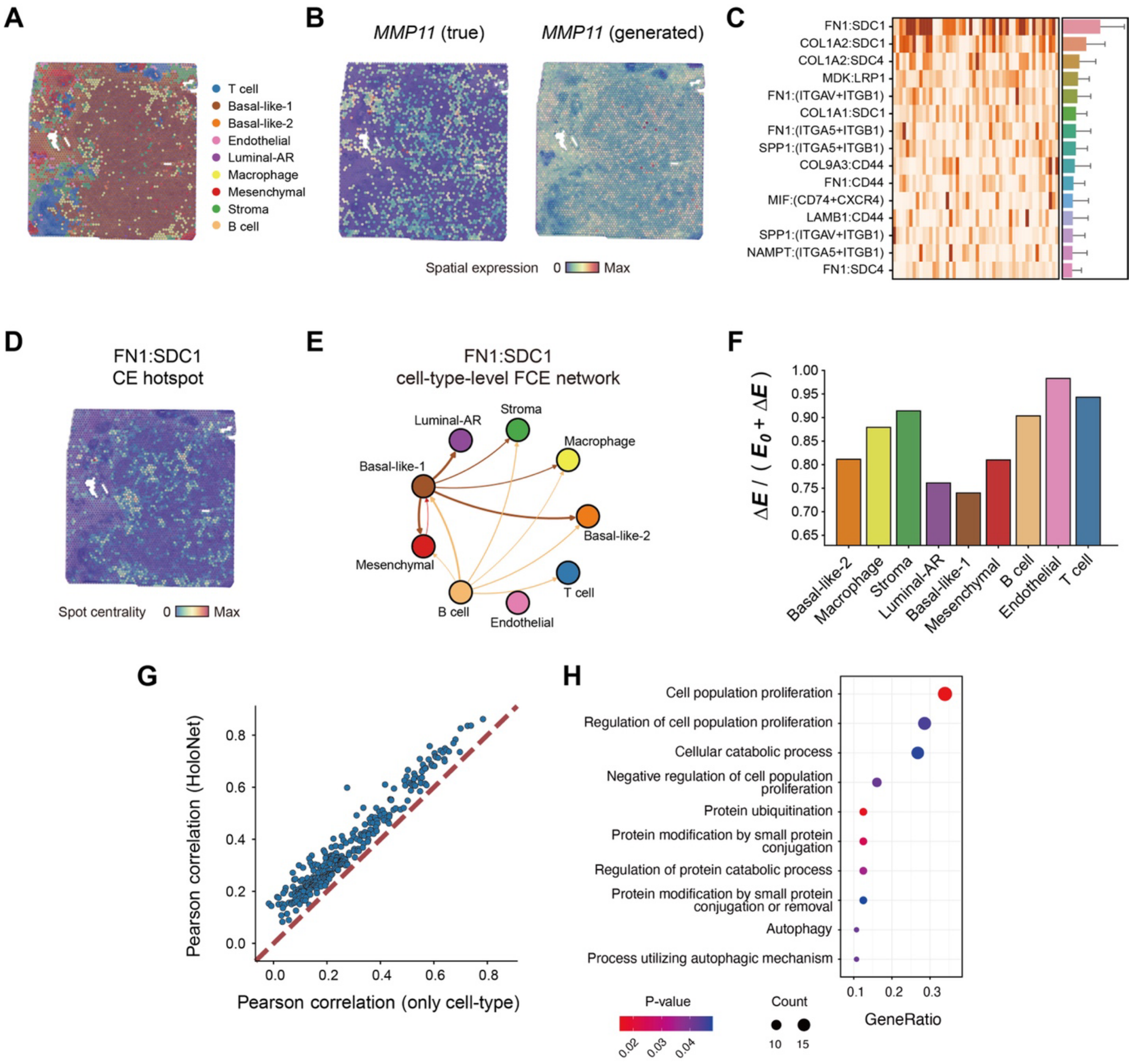


**Figure S7. Application of HoloNet to breast cancer Visium dataset from Wu et al.[22].**

**(A)** The cell type with the highest percentage in each spot. **(B)** The true (left) and HoloNet-generated (right) expression profile of *MMP11*. **(C)** Top 15 ligand–receptor pairs (138 pairs in total) with the highest view attention weights. Heatmap displays the attention weights of each view obtained from repeated training for 50 times. Bar plot depicts the mean values of the attention weights of each view, and the error bars represent standard deviations **(D)** The CE hotspot plot of FN1:SDC1. **(E)** Cell-type-level FN1:SDC1 FCE network for *MMP11*, similar as **Figure 3E**. **(F)** The proportion of the expression change caused by CEs ($\Delta E$) of the sum of $\Delta E$ and the baseline *MMP11* expression ($E_0$) in each cell-type. **(G)** Performance comparison between HoloNet and the models only using cell-type information. The comparison is based on tasks for predicting the expression level of each gene. **(H)** Gene ontology (GO) enrichment for the top 50 genes with relatively higher performance improvements after considering CEs, with all 285 genes as the background genes.

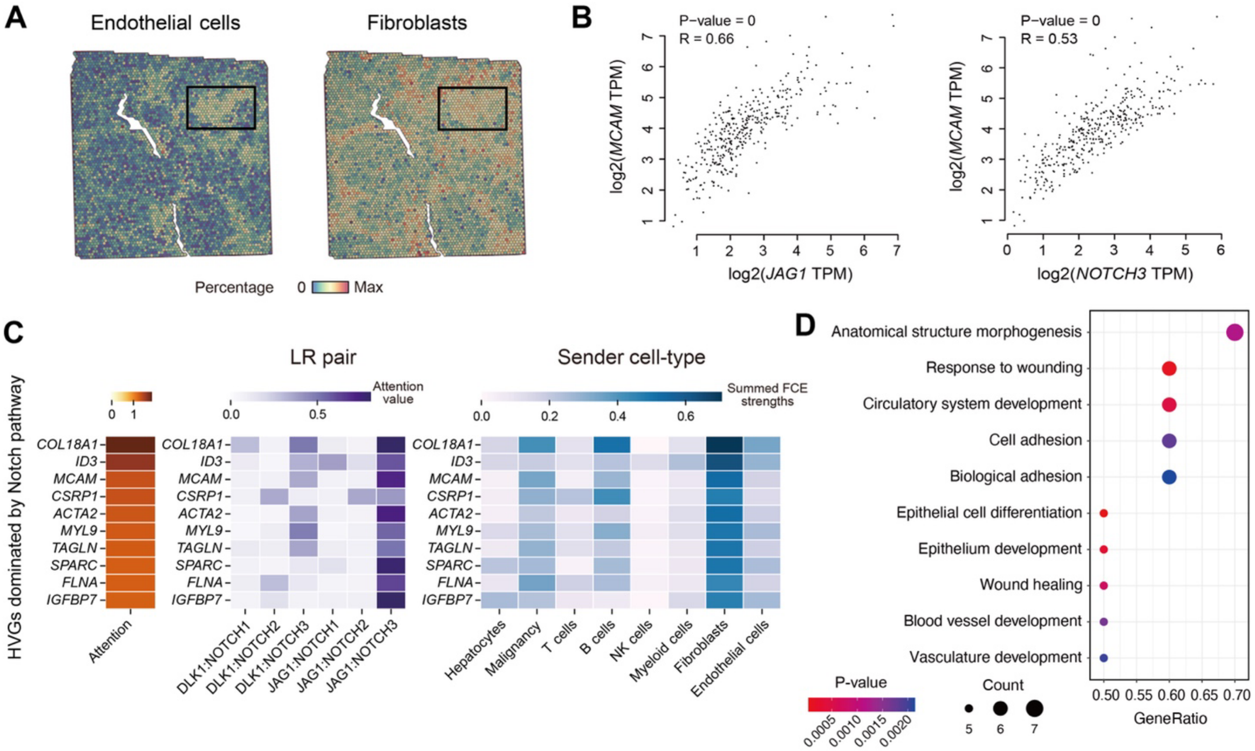


**Figure S8. Application of HoloNet to liver cancer dataset.**

**(A)** The percentages of endothelial and fibroblasts in each spot. The black box indicates the area where the endothelial cells aggregate, and the area also is the possible location of blood vessels. **(B)** Correlation analysis between *JAG1* and *MCAM*, and between *NOTCH3* and *MCAM* expression based on TCGA liver tumor samples. **(C)** Heatmaps for NOTCH pathway related genes, and which ligand receptors affect these genes, and these FCEs come from which cell types. **(D)** GO enrichment for the 10 genes related to NOTCH pathway.

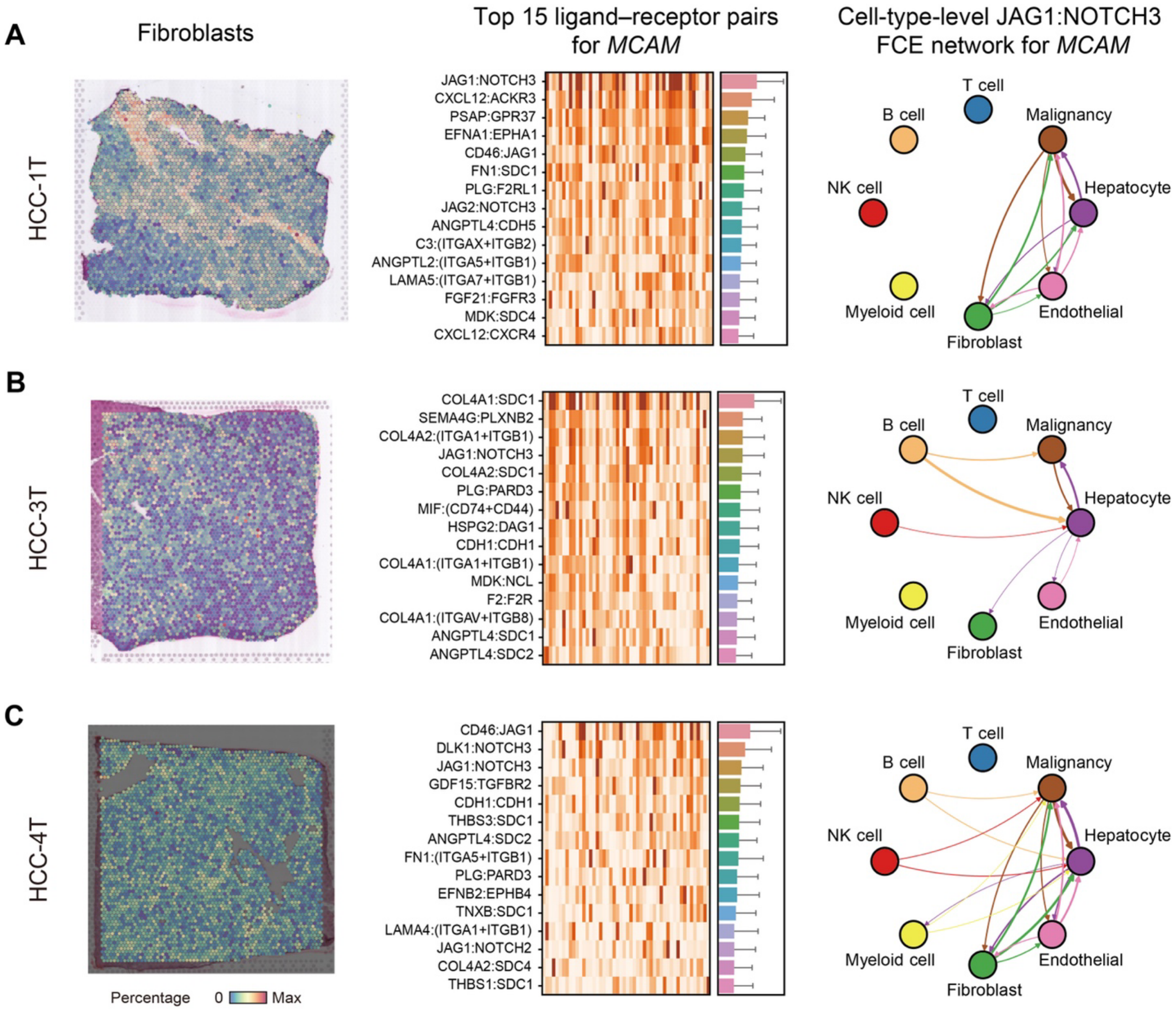


**Figure S9. Application of HoloNet to HCC-1T, HCC-2T and HCC-4T.**

**(A-C)** Displaying the results of HoloNet in the similar way as **Figure S8A, 5E and 5G**, to verify the findings based on HCC-2T.

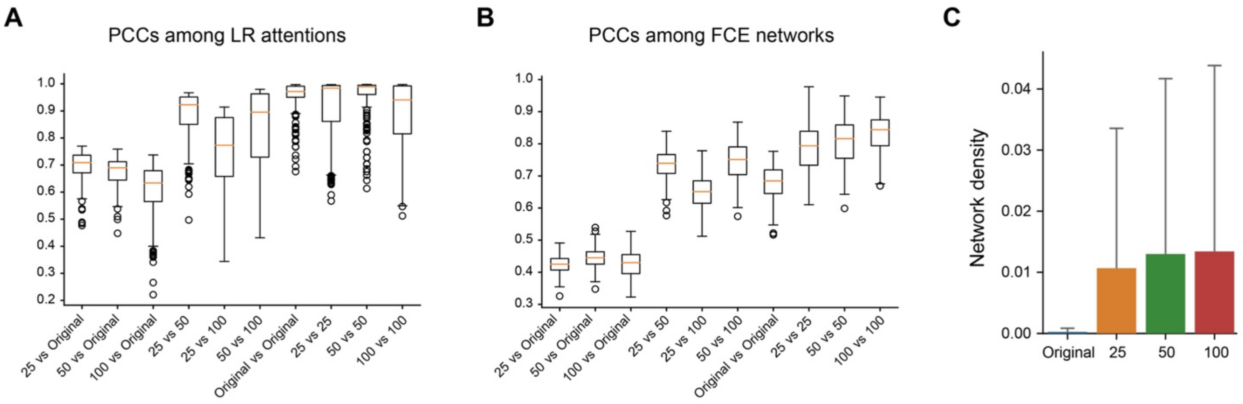


**Figure S10. Validation the stability of HoloNet in the mouse hippocampus Slideseq-v2 dataset.**

**(A)** Pearson's correlation coefficients (PCCs) among LR attentions for each target genes. The mean value of each column is in **Figure 6B**. Taking 'Original vs Original' as an example, a target gene can obtain a LR attention list. We calculated Pearson's correlation coefficient (PCC) of the attention lists among 3 repeatedly down-sampled dataset. This gets 3 correlation coefficients, and the mean value of 3 correlation coefficients is a point of 'Original vs Original' columns. There are 338 points in the columns corresponding to 338 target genes. **(B)** PCCs among FCE networks for each target genes. Similar to **A**. **(C)** Network densities of 5000-spot down-sampled datasets, and 5000-spot datasets after fusing 10μm spots within 25, 50, and 100μm.
